# Revisiting the Central Dogma: the distinct roles of genome, methylation, transcription, and translation on protein expression in *Arabidopsis thaliana*

**DOI:** 10.1101/2025.01.08.631880

**Authors:** Ziming Zhong, Mark Bailey, Yong-In Kim, Nazanin P. Afsharyan, Briony Parker, Louise Arathoon, Xiaowei Li, Chelsea A. Rundle, Andrew Behrens, Danny Nedialkova, Gancho Slavov, Keywan Hassani-Pak, Kathryn S. Lilley, Frederica L. Theodoulou, Richard Mott

## Abstract

**Background:** We investigated the flow of information from genome sequence to protein expression implied by the Central Dogma, to determine the impact of intermediate genomic levels in plants.

**Results:** We performed genomic profiling of rosettes in two *Arabidopsis* accessions, Col-0 and Can-0, and assembled their genomes using long reads and chromatin interaction data. We measured gene and protein expression in biological replicates grown in a controlled environment, also measuring CpG methylation, ribosome-associated transcript levels and tRNA abundance. Each omic level is highly reproducible between biological replicates and between accessions despite their 0.5% sequence divergence; the single best predictor of any level in one accession is the corresponding level in the other. Within each accession, gene codon frequencies accurately model both mRNA and protein expression. The effects of a codon on mRNA and protein expression are highly correlated but are unrelated to genome-wide codon frequencies or to tRNA levels which instead match genome-wide amino acid frequencies. Ribosome-associated transcripts closely track mRNA levels.

**Conclusions:** In the absence of environmental perturbation, neither methylation, tRNA nor ribosome-associated transcript levels add appreciable information about constitutive protein abundance beyond that in DNA codon frequencies and mRNA expression levels. The impact of constitutive gbM is mostly explained by gene codon composition. tRNA abundance tracks overall amino acid demand. However, genetic differences between accessions associate with differential gbM by inflating differential expression variation. Our data show that the Central Dogma holds only if both sequence and abundance information in mRNA are considered.

## Background

Numerous studies in plants, fungi and animals have shown that the relationship between protein and mRNA expression levels is only moderate, with correlations typically around 0.5-0.6 [1–3]. This phenomenon is thought to be a consequence of several factors, principally, different rates of synthesis and degradation of mRNA and proteins [4], compounded with buffering and cross-talk between different spatial and temporal contexts of expression, protein length [5], measurement bias and inaccuracy [6] and, potentially, how the data are processed.

Recent advances in genomics technologies have made it possible to assemble genomes almost perfectly, to quantify DNA methylation and other epigenetic marks, and to measure protein and transcript expression accurately at scale. By integrating these data across omic levels we can now test if the flow of information supports the direction of the Central Dogma from lower to higher levels, namely genome ⇒ epigenome ⇒ transcriptome ⇒ proteome. (Here we have added epigenetics to the Central Dogma’s flow between genome and transcriptome; the genomic impacts of the environment are assumed to act via the epigenome). Modelling between different omic levels for the same gene addresses the question of correlations, while modelling across genes within each level reveals which factors act differentially between genes.

If we use lower omic levels (primary DNA sequence features and epigenetic marks) to predict higher levels (transcriptome, proteome), and ultimately phenotype [7], then three questions are particularly relevant: First, which features of the underlying DNA sequence are most predictive of higher omic levels? Second, how much of the information about protein expression levels encoded in the basal genome sequence is mediated through intermediate epigenetic and transcriptomic levels, and does it pass through multiple causal pathways? Third, in plants, what is the role of constitutive gene body CpG methylation (gbM) in controlling gene and protein expression? This role has been debated, and indeed it has been suggested that constitutive gbM might have no function, although it appears to be evolutionarily conserved [8], under selection [9] and to play a role in adaptation [10]. Furthermore, is it unclear how constitutive gbM in a specific gene is established [11, 12].

Large-scale population-based studies can reveal how genetic and environmental variation impact expression of different omic levels, but there is an equally strong case for looking in detail across many biological replicates at smaller systems where genotype and environment are tightly controlled and in which each omic level is measured in biological replicates as reproducibly as possible. Here, we employed the second approach. We performed detailed genomic profiling of two *Arabidopsis thaliana* accessions: Col-0 (the reference) and Can-0. The latter accession originates from the Canary Islands and is phenotypically adapted to an environment quite different from the central European origin of Col-0 [13]. Can-0 has about double the number of sequence differences from Col-0 compared to other accessions [14], and has been variously characterised as a relict by [15] and as “admixed” and distinct from four main genetic clusters (Europe, Madeira, Asia, Africa) identified by long-read sequencing of 70 accessions in [16]. Can-0 and Col-0 therefore represent genetically distinct lineages of Arabidopsis, so identifying shared characteristics between these accessions in contrast to those that differ may reveal answers to some of the questions described above.

We re-assembled the accessions’ genomes and measured constitutive CpG gbM for each gene. We re-annotated each assembled genome to produce accurate data across omic levels, aiming to eliminate reference bias. To try to eliminate environmental perturbations, we grew multiple biological replicates of each accession under the same climate-controlled, long-day environment in growth chambers. We quantified mRNA, tRNA, ribosome-associated transcripts, and protein abundances in rosette leaves of defined developmental age.

These data enabled us to explore how variation in genome, epigenome, transcriptome, and proteome are related. We first tested whether lower omic levels predict higher levels, and whether the information they encode was unique to that level, using the fraction of variation across genes in a focal omic level that is explained by variation in lower levels for this purpose. We asked if the influence of gbM on higher omic levels is subsumed by the information encoded by genome sequence, specifically in gene codon frequencies, if these codon frequencies affect mRNA and protein expression in similar ways, whether the impact of each type of codon is related to its genome-wide frequency, and how the levels of tRNAs relate to mRNA and protein expression. Finally, we examined differences in methylation and expression between Col-0 and Can-0 to test whether the former are related to the latter, which types of differences are most important, and what this tells us about the causal effects of methylation. Our analysis uncovers some unexpected yet important relationships, whilst showing others are insignificant under the experimental conditions employed here.

## Results

### Col-0 and Can-0 genome assembly and annotation

We produced high-quality *de novo* assemblies of the Col-0 and Can-0 genomes, from a combination of long (HiFi, ONT) and short (Illumina) reads, using Omni-C chromatin interaction data to confirm our assemblies and orient scaffolds (**Supplementary Table 1, Figure 1A**). Seventeen and five gaps remain in our assemblies of Col-0 and Can-0 respectively, all concentrated in rDNA arrays and centromeres (**Figure 1B**). However, we obtained excessively high depth of coverage of both ONT and HiFi reads in these repeat regions, suggesting the numbers of tandem repeats in centromeres may be underestimated and, potentially, variable between nuclei (**Figure 1A**). Outside of large tandem repeats, Omni-C data indicate that the assemblies are structurally accurate (as shown by the Pretext contact maps in **Supplemental Figures S1** (Col-0) and **S2** (Can-0), as do the BUSCO and QV gene content statistics (in **Supplemental Table S1**).

**Figure 1.**
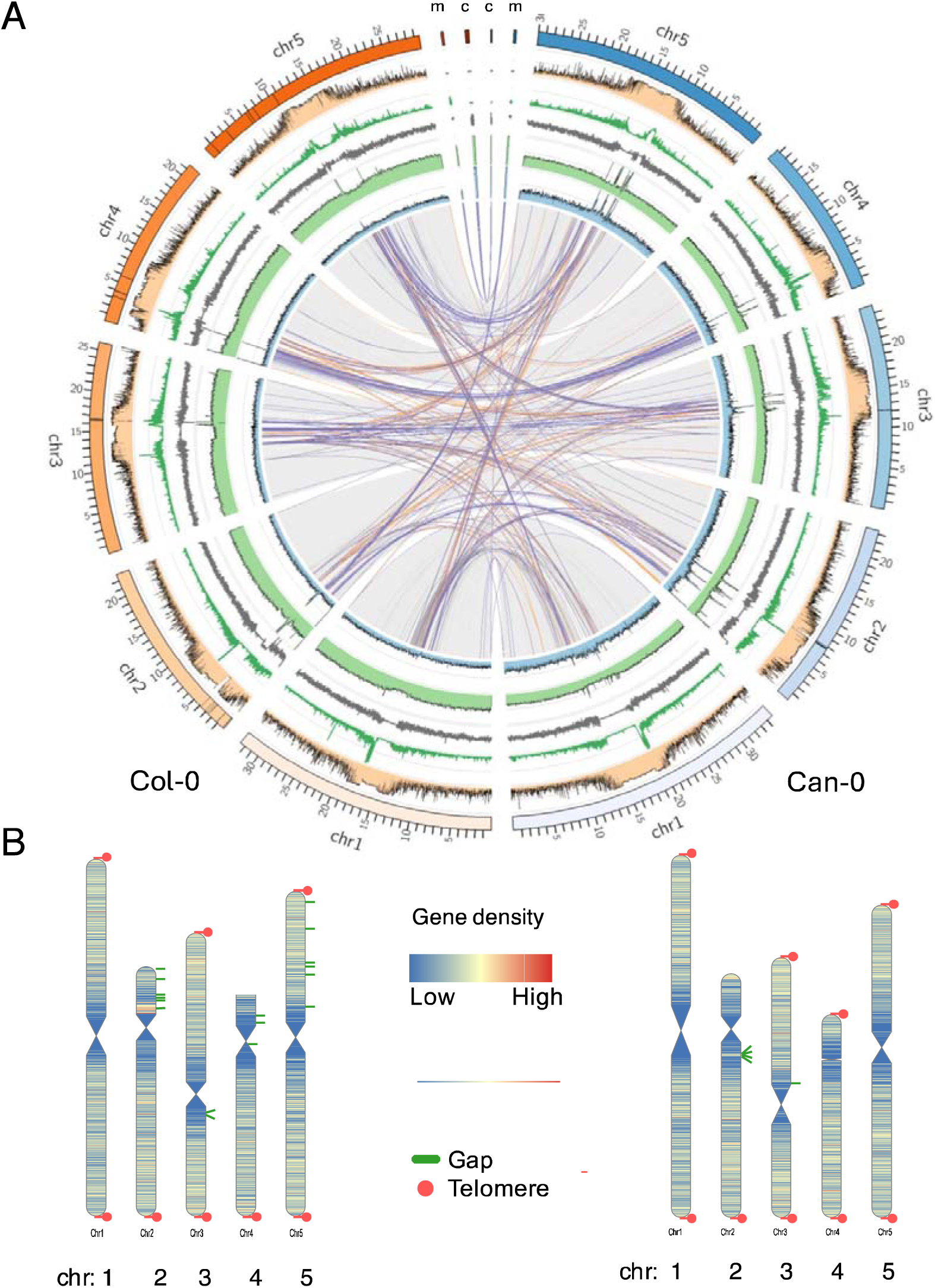
Assembly of Col-0 and Can-0 genomes. (A) Circos plot comparing the Col-0 (orange set, left) and Can-0 (blue set, right) genomes. Links in the middle show genomic rearrangements. Purple links: inversions, orange links: translocations between different chromosomes. Light blue: HiFi coverage, light green: ONT coverage. Grey line panel: GC content of the genome. Dark green line: repeat density, Orange filled panel: percent CpG methylation. (B) Cartoons of chromosome (chr) assemblies of Col-0 (left) and Can-0 (right) showing gene densities, the positions of assembly gaps (green) and indicating where the assemblies reached into the telomeres (red dots).

We aligned our Col-0 sequence with five published Col-0 assemblies, namely the old reference TAIR10 [13], Col-CEN [17], Col-XJTU [18], Col-Rian [16], and the new community reference Col-CC (Genbank reference GCA_028009825.2). The numbers of differences are shown in **Figure 2** and are generally small, e.g. there are only about 10,000 SNP differences between the Col-0 assemblies, and almost all of the observed differences are in the numbers of tandem repeats. Some differences are likely to be artefacts from different assembly algorithms; indeed, our Col-0 most closely resembles the Col-XJTU assembly, where the same software and similar pipelines were employed (**Supplemental Table S2).** Comparison of Col-CC and Col-Cen shows similar numbers of differences, so our Col-0 assembly is not unusual.

**Figure 2.**
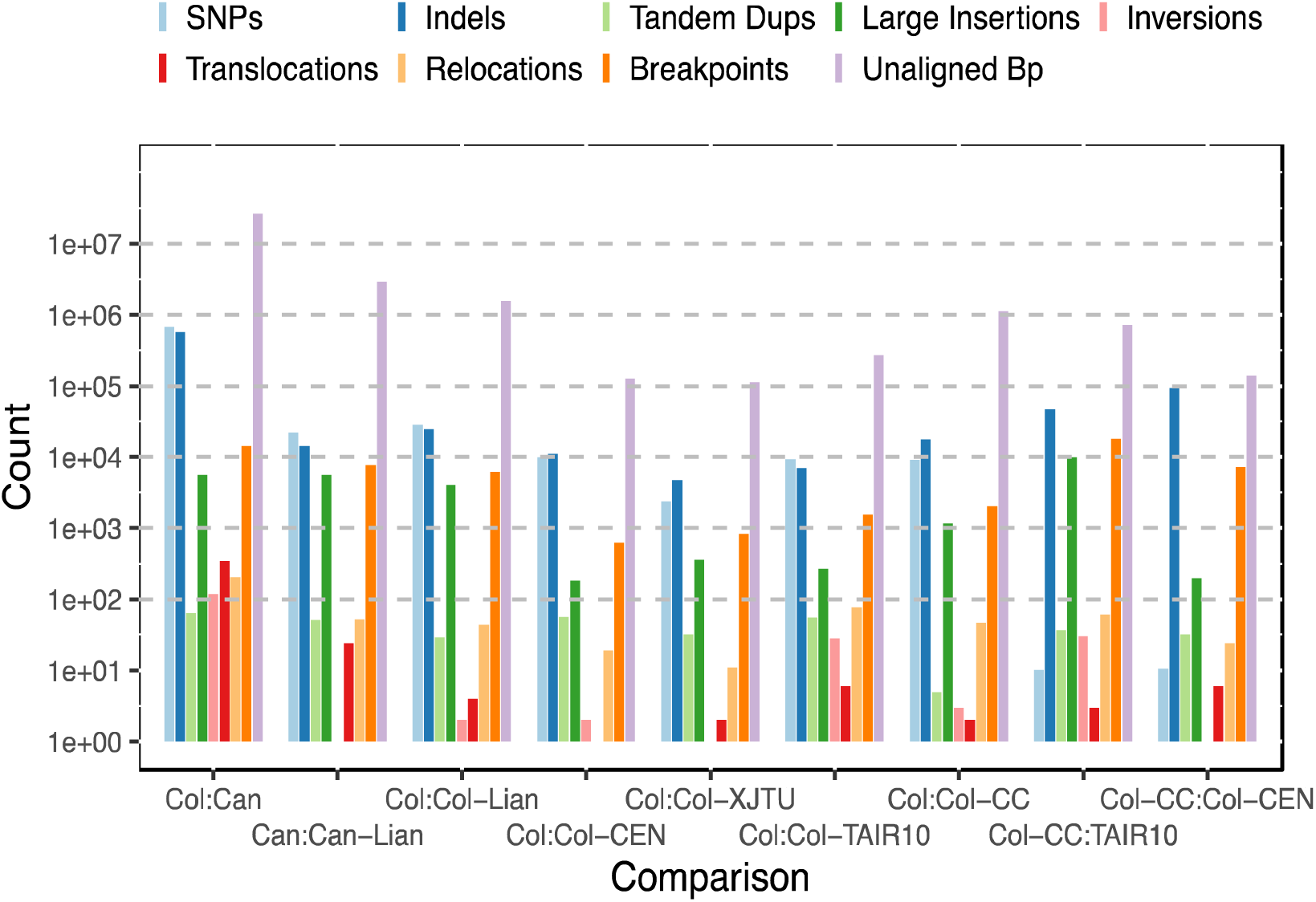
Counts of differences as computed by dna_diff [20] between our Can-0 (blue background) and Col-0 (yellow background) assemblies and with four other Col-0 and one other Can-0 assemblies, namely Can-Lian, Col-Lian:[16], Col-CEN [17], Col-XJTU [18], Col-TAIR, the TAIR10 reference, Col-CC: the community consensus assembly (Genbank id GCA_028009825.2). Pink background shows comparisons between selected other Col-0 assemblies.

We also compared our Can-0 assembly to that reported in [16] (named Can-Rian here) and again found similar numbers and patterns of differences as observed between different long-read Col-0 assemblies (**Figure 2**, (**Supplemental Table S2**). We conclude that all these assemblies have similar accuracies and that most of the differences occur within highly repetitive regions and represent in part algorithmic artefact. However, we cannot exclude the possibility of small numbers of genuine sequence differences in the germplasm used as a sequencing substrate, arising in unstable tandem repeat regions.

Henceforth, "Col-0” and “Can-0” refer to our assemblies and annotations of these accessions, unless otherwise stated. Where we calculate the same statistic in Col-0 and Can-0, the numbers are reported as an ordered pair. For example, our assemblies’ lengths (Col-0: 133.23 Mb, Can-0: 133.09 Mb) and N50 values (18.4 Mb, 12.5 Mb) are very similar to those obtained in [17, 18].

We annotated both genomes, using *ab initio* gene prediction applied to both short-read (Illumina) and long-read (Iso-Seq) RNAseq data from rosette leaves sampled at the 9-leaf stage. We annotated 28,763 and 28,532 protein-coding genes and 47,596 and 45,756 alternatively spliced isoforms (**Supplemental Table S3**). These counts include duplicated genes within each accession. If we exclude duplicates then there are 24,325 pairs of orthologous genes between Col-0 and Can-0 of which 23,081 are also annotated in Araport11 [19]. Several hundred genes are unique to each annotation, as shown in the Venn diagram **in Figure 3**. **Supplemental Files S1, S2** contain the annotations of the genomes as GFF files. **Supplemental Files S3 and S4** contain the mRNA and amino acid sequences of the Col-0 predicted genes and **Supplemental Files S5 and S6** contain those for Can-0.

**Figure 3.**
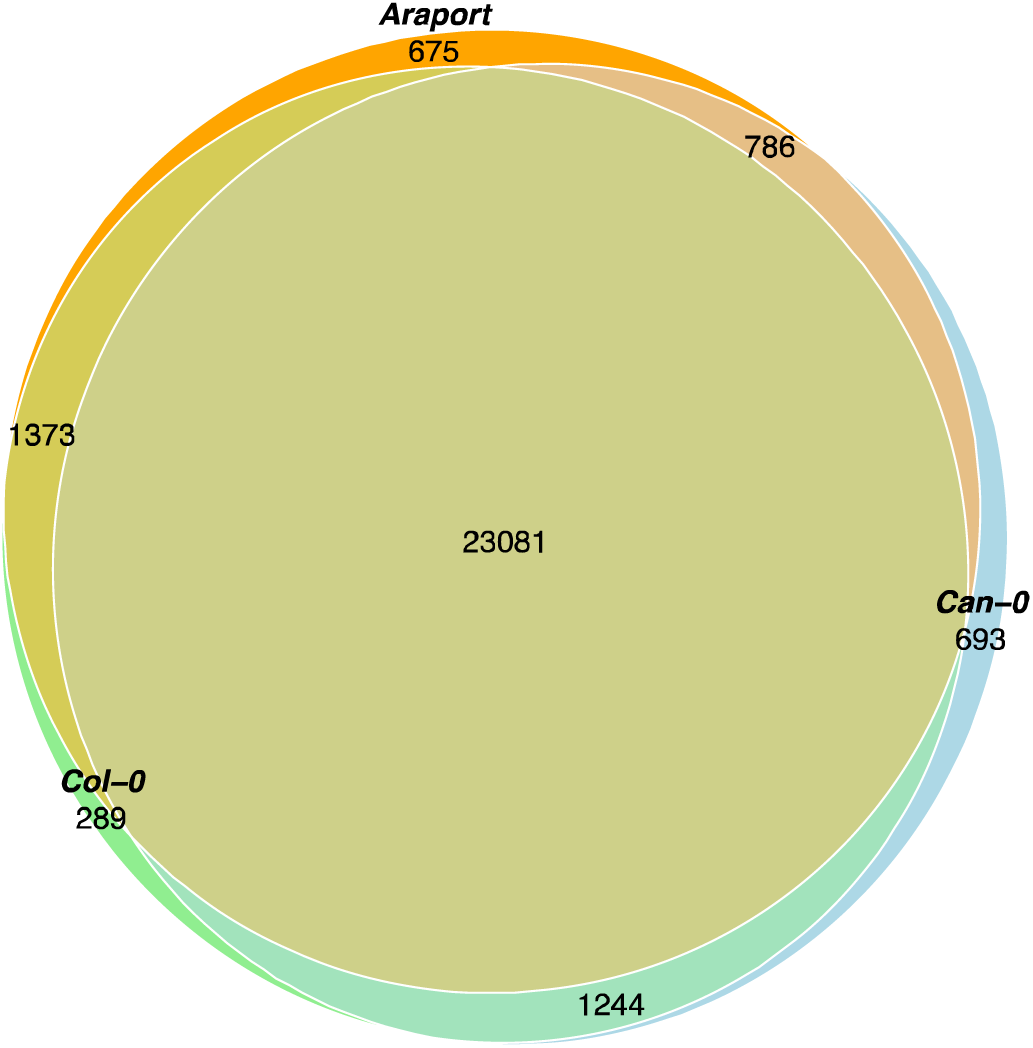
Venn diagram showing overlaps between genes annotated in Col-0, Can-a0nd Araport11 [19]. The numbers shown are the counts of genes in the different intersections. For example, there are 289 genes only found in the Col-0 annotation, while 1,373 genes are common to Col-0 and Araport but absent from Can-0.

Taking the primary isoform in each accession, in total 20.30% of orthologous coding sequences are identical at the amino acid level, and 52.67% differ at no more than six codons (see below). However, 6.96% differ by more than 100 codons. Since 35.09% of homologous gene pairs have different numbers of annotated isoforms, there is some ambiguity in these statistics.

### mRNA and protein expression levels are highly reproducible but relatively poorly correlated

We quantified the abundance of mRNAs, ribosome-associated RNAs, and proteins in Col-0 and Can-0 rosette leaves raised in growth chambers under identical long day conditions and harvested at the same 9-leaf developmental stage (**Supplemental Table S4**). By minimising environmental and temporal variation we thereby focused on internal sources of variation in gene and protein expression and ribosome association. We applied quality control filters to remove duplicated genes, genes longer than 6 kb, and genes without a 1-1 ortholog between the two accessions, leaving 22,606 genes.

Of these genes, 18,226 were expressed in Col-0 (and 772 only in Col-0) and 18,200 expressed in Can-0 (746 only in Can-0). There were 17,454 ortholog pairs with mRNA expression in at least one replicate in both accessions. After removing genes with premature stop codons, 17,414 expressed genes remained for analysis. If we further only retain those genes with expressed mRNA in all five biological replicates in each accession then there are 15,669 in Col-0, 15,669 in Can-0 and 15,215 in both. Our downstream analyses do not employ this additional filter except where noted.

Proteins were analysed using a label-free data-independent acquisition (DIA) workflow, employing intensity-based absolute quantification (iBAQ; [64]) to derive comparative protein abundance from mass spectrometry data between the two accessions. This enabled quantification of 8,915 proteins common to Col-0 and Can-0. Both mRNA and protein expression were detected in both accessions for 7,771 ortholog pairs, whilst a further 9,633 pairs had mRNA expression in both accessions but no observed protein expression. To some extent, this reflects the comparatively lower coverage of proteomic data, but it also suggestive of extensive post-transcriptional control. Interestingly, 28 genes had protein but no mRNA expression in Col-0 and 32 in Can-0, perhaps indicating RNA instability. If we only retain genes expressed in all replicates, then there are 7,721 genes expressed in mRNA and protein in both accessions and 7,494 expressed in mRNA but not protein in both accessions; no genes were expressed in protein but not mRNA.

Both mRNA and protein measures were highly reproducible across replicates **(Supplemental Table S5**); for genes with both mRNA and protein expression, correlations of log-transformed mRNA levels between 5 replicates within an accession all exceed 0.98, as did correlations between 4 replicate protein levels within an accession (all correlations are between log-transformed data unless otherwise stated). Correlations between accessions are also very high; all mRNA replicates exceed 0.95, despite the presence of many differentially expressed genes (discussed later). The strength of these mRNA correlations was slightly lower for genes without protein, although all correlations between replicates still exceeded 0.94. Correlation of protein expression between each accession always exceeded 0.91 among replicates.

We then combined the expression levels across replicates for each gene by taking geometric means. Looking across genes within each accession, the most abundantly expressed genes and proteins greatly exceed the respective median level: 2,151-fold, Col-0; 1,370-fold, Can-0 for mRNA and 640-fold; 591-fold, respectively for protein (**Figure 4 A,B**). We report fold changes relative to the median rather than the full dynamic range of expression because the latter is strongly biased by genes expressed at near-zero levels. Protein dynamic range is well established to exceed that of RNA but is not generally captured in proteomics workflows, and especially not in tissues such as leaves which are dominated by a small number of highly abundant proteins [21, 22]. Most of the highly expressed genes and proteins are involved in photosynthesis, as would be expected for leaf tissue.

**Figure 4.**
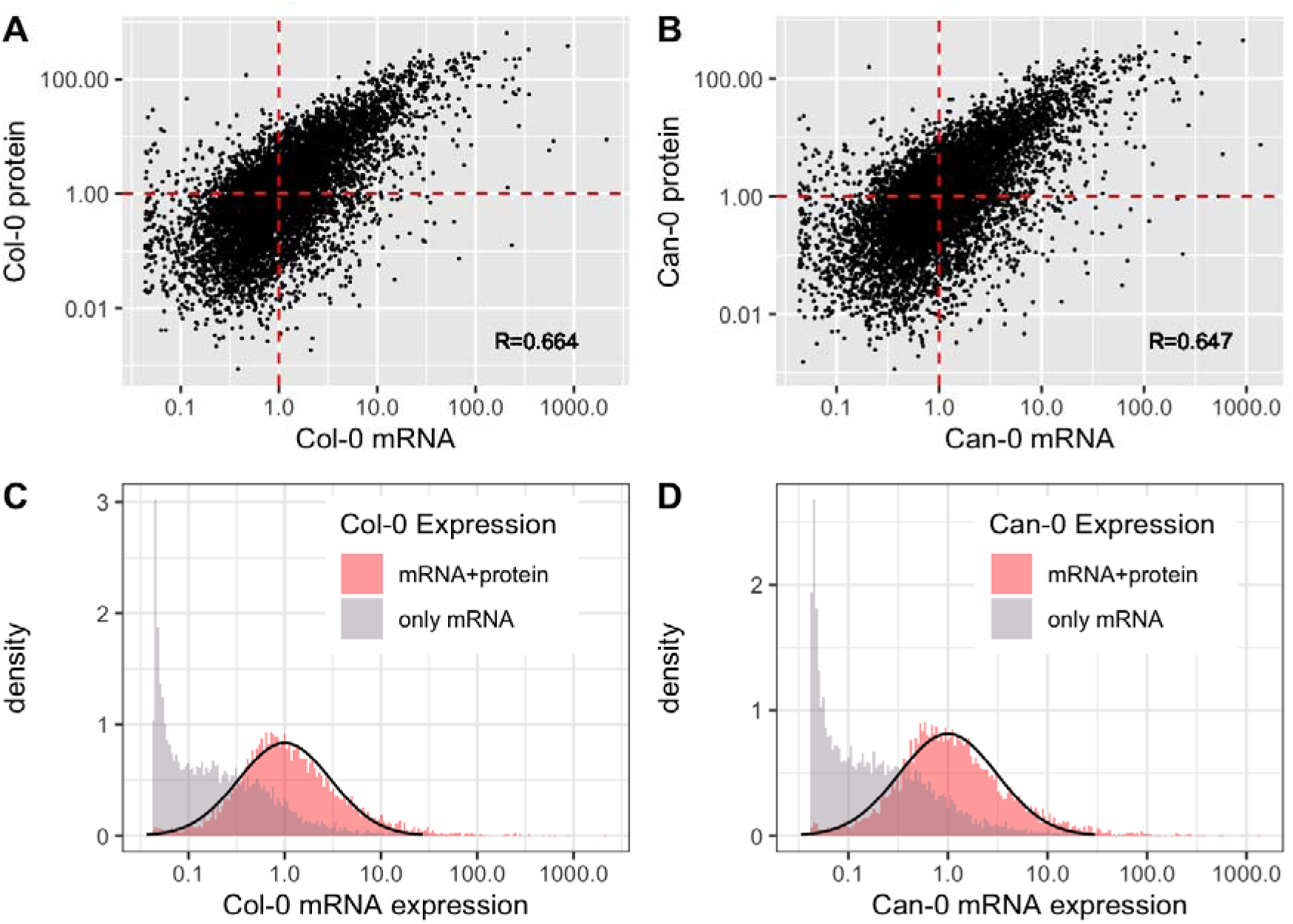
**The** distribution of mRNA and protein expression, scaled so that the median level of expression of genes with both protein and mRNA expression is equal to 1. A, B: scatter plots of mRNA (x-axis) vs protein (y-axis) expression for orthologous genes in Col-0 (A) and Can-0 (B). Dotted red lines show medians. C,D: Histograms of mRNA expression for genes with (pale red) or without (grey) detectable protein expression in Col-0 (C) and Can-0 (D). The black curves indicate lognormal densities fitted to the mRNA+protein histograms using robust estimates of mean and standard deviation. Expression scales are logarithmic throughout.

To analyse correlations within and between mRNA and protein expression we focussed on the 7,771 ortholog pairs of genes with mRNA and protein expression in both Col-0 and Can-0. After log-transformation the correlation between mRNA and protein within an accession was (0.664, 0.647), about 50% higher than without log transformation (**Table 1**, **Figure 4A,B**), and similar to that reported in maize leaves [1]. Despite differences in measurement and analysis methodologies the most accurate predictor of protein expression is not mRNA from the same accession but rather protein from the other accession **(Table 1).** We return to this point below.

**Table 1.** Pearson correlations between mean expression levels of mRNA and protein in Col-0 and Can-0 across 7,771 genes. Blue background: correlations between raw expression values. Orange background: correlations between log-transformed values, using the transformation y=log _10_(x+1) i.e. adding a pseudo count of 1 unit.

|  | mRNA<br>Col | mRNA<br>Can | protein<br>Col | protein<br>Can |
| --- | --- | --- | --- | --- |
| mRNA Col | 1 | 0.9722 | 0.6635 | 0.6382 |
| mRNA Can | 0.9679 | 1 | 0.6444 | 0.6471 |
| protein Col | 0.3101 | 0.3960 | 1 | 0.9154 |
| protein Can | 0.3104 | 0.4075 | 0.9406 | 1 |

The distributions of mRNA gene expression were markedly different depending on whether protein expression is also observed or not (**Figure 4 C,D)**, with the former following an approximately lognormal distribution. Genes without detectable protein expression have a complex mRNA distribution, comprising a spike of genes with very low expression and a shoulder of intermediate expression, with a thin tail of highly expressed genes. The spike of genes with near-zero expression is due to those genes not expressed in all replicates. **Supplemental Figure S3** shows the same distributions in Figure 4 omitting the 9,633 - 7,494 = 2,139 imperfectly replicated genes.

### Ribosome associated transcript levels closely resemble standard mRNA expression for genes with protein expression

We next asked if transcripts associated with ribosomes were better correlated with protein expression. Ribosome-associated RNAs were quantified in six biological replicates each of Col-0 and Can-0 rosettes using 3’Ribo-seq [23] (expression levels are in **Supplemental File S4**). In total 17,513 of their 1-1 orthologs had detectable ribosome-associated transcripts (ribo-mRNA hereafter) in both accessions. Within the 7,771 genes with protein expression, ribosome-associated transcripts behaved very similarly to the mRNA data described above; 7,620 (97.9%) genes were also associated with ribosomes, and the correlation of log-transformed ribo-mRNA and mRNA expression levels was 0.9530 (Col-0), 0.9574(Can-0). Correlation among the six biological replicates always exceeded 0.97 within an accession and exceeded 0.94. between accessions (**Supplemental Table S5**). Their correlation with protein expression was (0.6490, 0.6415), very similar to that observed for mRNA determined by RNA-seq.

Within the 9,673 genes for which mRNA but no protein was quantified, the pattern is slightly different. Of these, 6812 (70.42%) also show ribosome association, and the correlation of expression levels between mRNA and ribo-mRNA among these genes remained high, at (0.9303, 0.9451). All correlations between replicates within an accession exceeded 0.89 but between accessions, the correlations were lower with a minimum of 0.67. Only 59 genes are expressed in ribo-mRNA but absent from mRNA. Among the 29.58% of genes without ribo-mRNA expression, average mRNA expression was reduced by factors of (5.6, 5.4). Thus, the spikes of genes with very low expression seen in **Figure 4 C,D** are absent among ribosome associated transcripts. Apart from this difference, the ribo-mRNA expression levels and patterns were essentially interchangeable with those of mRNA, and so for the remainder of this study we use the mRNA expression data.

### Constitutive CpG methylation is reproducible across assay type and between accessions

We measured CpG methylation using both bisulphite-converted Illumina reads and ONT long reads, the latter collected as a by-product of generating sequence for *de novo* genome assembly. Because of the larger amount of leaf tissue required, plants for ONT sequencing were grown under short day conditions, whereas long day conditions were used for bisulphite sequencing to enable direct comparisons with the RNA-seq and proteomics datasets. We quantified each methylated CpG dinucleotide as the percentage of methylated bases from reads covering that CpG position. As **Figure 1A** (orange track) shows, uniformly high levels (>75%) of CpG methylation occur throughout the centromeres but methylation is variable in other regions of chromosomes. Across CpG sites, the correlation between bisulphite and ONT gbM values was 0.94 in both Col-0 (2.6 M sites) and in Can-0, (2.8 M) despite the different growth conditions (**Supplemental Figure S4)**. As the coverage of ONT data is superior to bisulphite sequencing and methylation readouts are less prone to GC bias [24] we therefore used ONT CpG methylation values in all further analyses.

We then computed gene-body methylation (gbM) for every annotated gene as the mean percentage methylation across all CpG dinucleotides within the genomic interval spanned by the gene, including exons and introns. We also quantified methylation in flanking regions around each gene for differential expression analysis (described later). The genome-wide distributions of gbM in Col-0 and Can-0 across gene expression categories (i.e., protein and mRNA, only mRNA, or no expression) are shown in **Figure 5**. All three distributions share a mode in gbM around 12%, but with differing upper tails. The correlation between gbM in Col-0 and Can-0 is slightly higher in genes with protein and mRNA detected (*R* = 0.890) than in genes with only mRNA (*R* = 0.830) but is surprisingly similar. **Figure 4** also plots the distribution of (6195, 6042) other genes without any expression at mRNA or protein. These contain subsets of (838, 952) genes where gbM exceeds 75%, and which are concentrated in the shoulders of the highly methylated centromeres of each chromosome, i.e. matching the local CpG methylation levels (**Figure 1**). However, most genes in all three categories have low levels of gbM under 25%; all modes are close to 10%. Thus, constitutive gbM varies only slightly across most genes. Below we discuss its impact on expression below, after considering codon composition effects.

**Figure 5.**
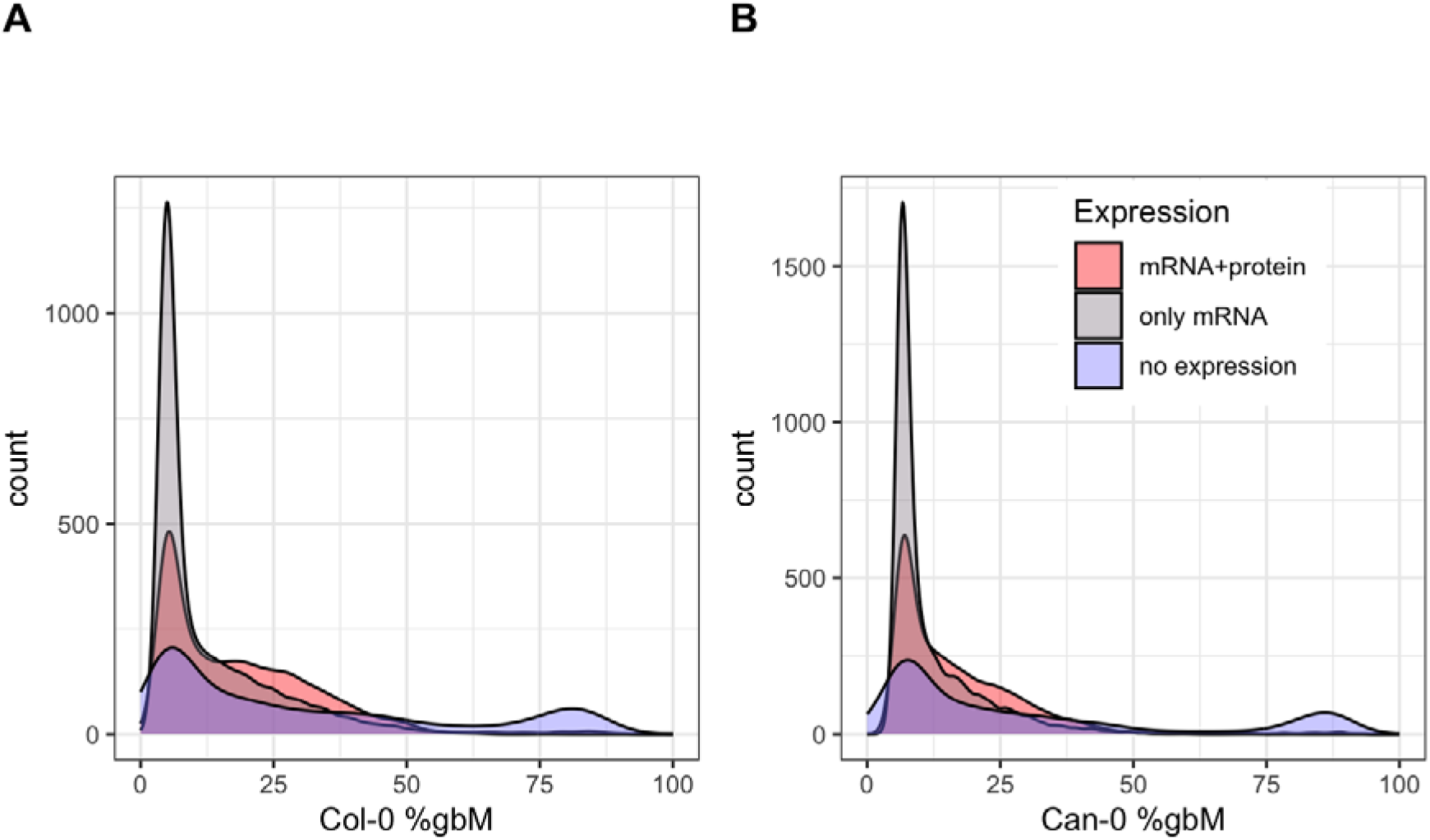
Distribution of gbM in (A) Col-0 and (B) Can-0 genes, categorised according to whether both protein and mRNA were detected or only mRNA detected, or not expressed.

### Codon composition affects expression

Codon composition is known to affect both mRNA and protein expression [25–27]. In our analysis we represented the coding sequence of each protein-coding gene by a vector of the 61 non-terminator codon frequencies, hereafter abbreviated to CDS. This representation therefore ignores the order of the codons. Among the 24,325 ortholog pairs of protein-coding genes, the median numbers of amino acids per gene in Col-0 and Can-0 are (349, 348) and the median difference in codon frequencies between orthologous genes in Col-0 and Can-0 (i.e. the sum of absolute differences in the counts of each of the 61 non-terminator codons) is 6 codons.

We modelled mRNA and protein expression of each gene in terms of these codon frequencies by fitting linear multiple regression models to the log-transformed expression levels, thereby estimating the expression effect (regression coefficient) of each codon. Under this model, if the codon *c* with multiple regression coefficient occurs times in gene, then its predicted log expression level is, where is the average expression level. Codons with positive codon effects increase expression and negative effects decrease it. The codon expression effects and their standard errors are shown in **Supplemental Table S7.**

We defined the standardised effect of each codon on expression as its expression effect divided by the standard error. Among 7,771 ortholog pairs with mRNA and protein expression in both accessions, standardised effects are highly correlated between mRNA and protein in both accessions (R=0.800, 0.800) (**Figure 6 A,B, Supplemental Figure S5**). These codon expression effects are highly reproducible between Col-0 and Can-0 (R=0.99 for both mRNA and protein). If we estimate mRNA effects by restricting attention to genes for which protein was not quantified, then the resulting mRNA codon effects are markedly less correlated with those modelled from genes with both protein and mRNA (, **Figure 6 C,D**).

**Figure 6.**
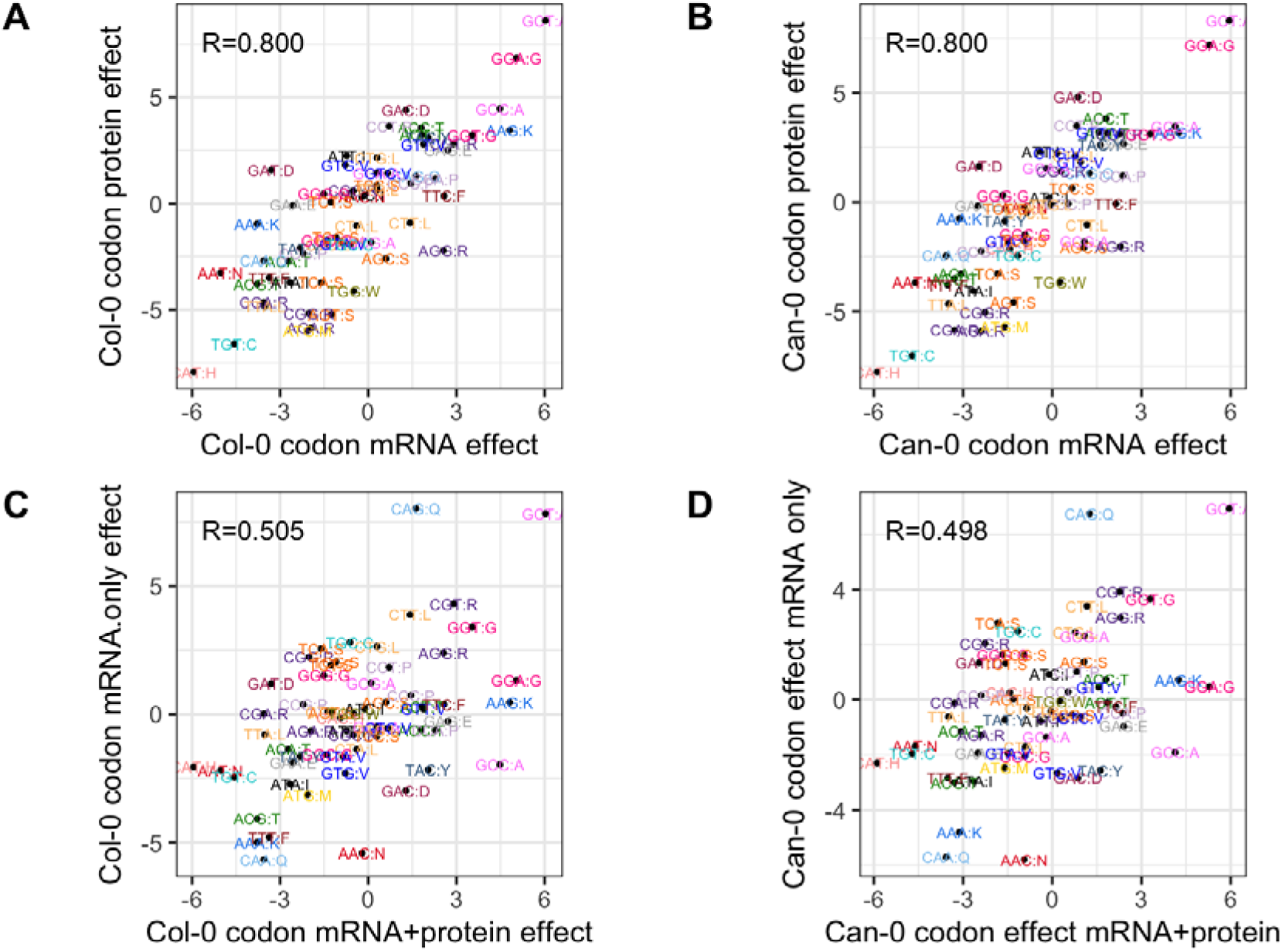
Codon effects on mRNA and protein expression. (A) Scatter plot of 61 codon-effects on log Col-0 mRNA expression (x-axis, represented as the T-statistic for each codon) vs the corresponding effects on log Col-0 protein expression. Each point is labelled with the codon and encoded amino acid, and all codons with the same amino acids share the same colour. Models were fitted to genes with both protein and mRNA expression in Col-0. (B) Similar analysis for Can-0. (C) Similar analysis comparing codon effects on mRNA expression in Col-0 estimated for genes with both mRNA and protein expression (x-axis) and for genes with only mRNA expression (y-axis). (D) Same plot as for (C) but in Can-0.

We next asked if these codon expression effects are related to global codon frequencies [28], to test the hypothesis that more frequent codons are associated with increased expression. We computed global codon proportions, either by summing the genomic gene codon frequences to give proportions independent of expression level, or by weighting the gene codon frequencies by mRNA or protein expression, thereby taking account of expression level. We then compared these proportions with the codon effects. The standardised codon effects for mRNA and protein expression are uncorrelated with overall codon abundance, defined as the relative fractions of codons across genes with both protein and mRNA expression (**Figure 7 A,B**; neither of the correlations of 0.065 and 0.235 are significant at P<0.05). Thus, more commonly used codons are not associated with higher gene or protein expression.

**Figure 7.**
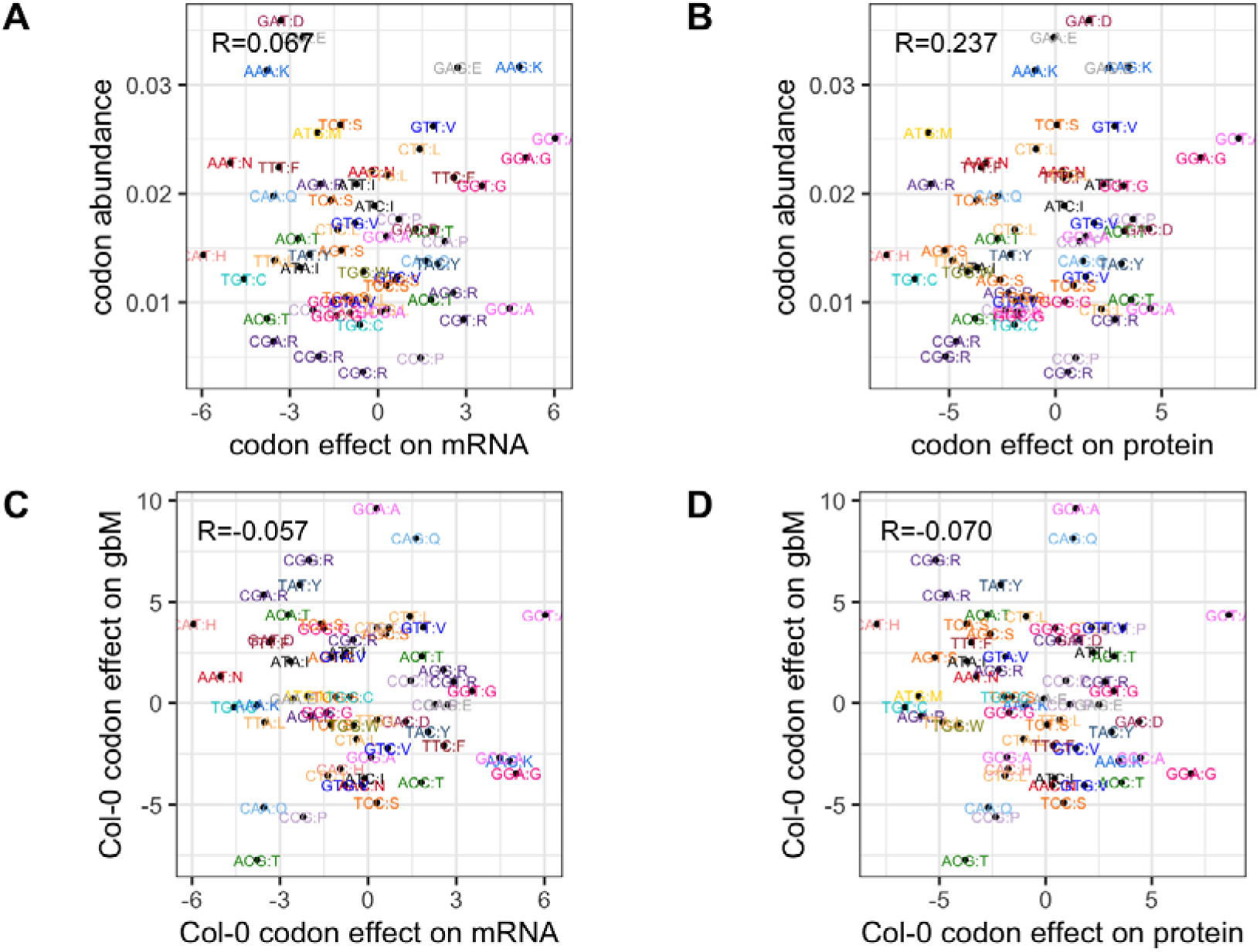
Lack of correlation between standardised codon effects in Col-0 and gbM and codon abundance. In each scatter plot, each point represents a codon and is colour-coded by the encoded amino-acid. A, B: standardised codon effects on gbM (y-axis) vs codon effects on mRNA expression (A) and protein expression (B). C, D: y-axis is codon abundance, defined as the fraction of codons in all genes with mRNA and protein expression, x-axis: codon effect on mRNA expression estimated multiple linear regression.

We also tested if codon frequencies correlate with gbM by fitting the same multiple regression models with gbM as the dependent variable in place of mRNA or protein expression. **(Figures 7 C,D)** show there is only weak correlation between the codon effects for expression and those for gbM. We return to the relationship between gbM and codons later.

### tRNA abundance tracks global amino acid frequency

We next asked if tRNA abundance was related to these codon expression effects and to codon abundance and gene expression. The genetic code is redundant, with 20 standard amino acids specified by 61 sense codons in eukaryotes. Isoacceptor tRNAs comprise families that accept the same amino acid, but which differ in their anticodon sequence, reflecting the fact that all amino acids other than methionine and tryptophan are specified by more than one codon. Isodecoder tRNAs carry the same anticodon but differ at their primary sequence at sites other than the anticodon. In common with other eukaryotes, Arabidopsis encodes tRNAs for only 45 sense codons plus the initiator tRNA-Methionine, the remainder employing third-base wobble base-pairing to effect translation [29–31].

We measured tRNA abundance using modification-induced misincorporation tRNA sequencing (mim-tRNAseq) [32, 33] in Col-0 and Can-0 leaves grown and harvested under the same conditions used to quantify mRNA and protein expression. Using the genomic tRNA database (GtRNAdb, [34]) annotation of Arabidopsis tRNA genes, we queried expression at 642 nuclear-encoded tRNA genes that also had Araport11 gene identifiers (**Supplemental Table S6)**, representing 224 distinct tRNA isodecoders. We observed non-negligible expression for 157 of these isodecoders. For each of the 46 anticodons, we calculated the relative expression across all isodecoders.

Non-organellar tRNA abundance levels are highly correlated between Col-0 and Can-0 (R=0.988; **Figure 8 A**). We found the relationship between codon frequencies and tRNA isodecoder abundance (**Figure 8 B,** *R* = 0.397) was obscured by the presence of codons without dedicated tRNAs (the vertically stacked codons on the left of **Figure 8 B**), although the figure also shows that some codons with no specific tRNA gene still have strong standardised effects. It is known which tRNAs translate these missing codons[31] (Supplemental Table S6a) and Figure 8C shows the scatter plot when we merge frequencies for codons which are translated by the same tRNAs; the correlation increases to *0.561* (**Figure 8 C**). Moreover, when we aggregate tRNA abundance and overall codon usage by the encoded amino acid (among genes with protein and mRNA expression) to produce isoacceptor frequencies, the correlation increases further *0.754* (**Figure 8 D;** results for Can-0 are very similar). The correlation reaches *0.811* when amino acid frequencies are weighted by protein abundance (**Figure 8 D**; a very similar relationship occurs when amino acid frequencies are weighted by mRNA abundance instead, because these are very highly correlated, (*R* = 0.979). The most discordant amino acid is tryptophan (W), which has higher tRNA expression than expected given its low frequency. Interestingly, tryptophan is the only amino acid apart from methionine encoded by a single codon. Additionally, it is encoded by UGG, which is in the same codon box as the UGA stop codon. High levels of its matching tRNA may be needed for it to compete with release factor, which may sample UGG codons.

**Figure 8.**
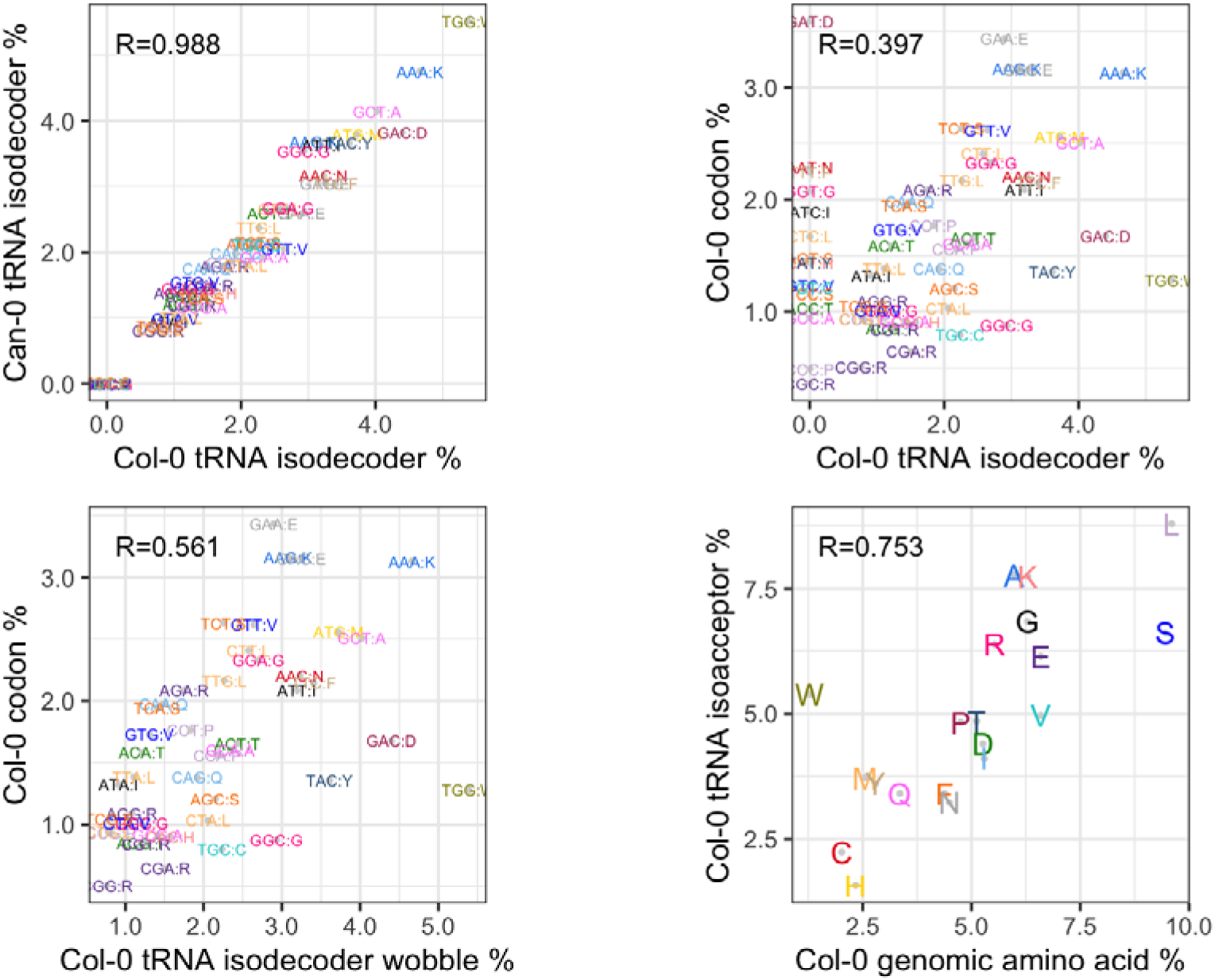
Relationships between tRNA abundance and codon and amino-acid frequencies. Codons and tRNAs are color-coded by encoded amino acid; the corresponding amino acid for each codon is specified in single letter code following a colon (i.e., in the format AAC:N; asparagine). Pearson correlation coefficients are shown in top left of each plot. (A) Col-0 tRNA isodecoder percentage (x-axis) vs Can-0 tRNA isodecoder percentage (y-axis). (B) Col-0 tRNA isodecoder percentage (x-axis) vs Col-0 codon frequency percentage. (C) Same as (B) except that codons without specific tRNAs are merged with the codons responsible for their translation to amino acids. (D) Col-0 codon fraction across all annotated genes (x-axis) vs Col-0 tRNA abundance aggregated by encoded amino acid (AA). Equivalent plots for Can-0 are very similar. If codon frequencies are re-weighted by the protein expression levels the plot are very similar with slightly higher correlations.

There is only a weak relationship between the tRNA abundances and our estimated codon effects on expression. The correlations between tRNA abundances and the mRNA codon effects are (0.286, 0.283, scatter plots in **Supplemental Figure S6 A, B**) and correlations with protein codon effects are (*0.249,0.234*, scatter plots in **Supplemental Figure S6 C, D).** These correlations are of borderline statistical significance (mRNA: *P* < *0.025*, protein: *P* < 0.057). In comparison, the correlation of *0.811* with amino acid frequencies satisfies *P* < 10^-14^. All codon-related statistics (regression coefficients and their standard errors, and tRNA abundance data) are in **Supplementary Table S7.**

### Modelling across omic levels reveals the importance of codon composition on expression and gene body methylation

We next asked which genomic features predict mRNA and protein expression levels across genes within each *Arabidopsis* accession. To model mRNA expression, the explanatory factors we considered were coding sequence DNA composition (referred to as CDS hereafter) and gene-body CpG DNA methylation (gbM, defined as the mean percentage of methylated CpGs within introns and exons). For protein gene expression we additionally considered mRNA expression as an explanatory factor. We also modelled gbM in terms of CDS. Taken together, these model choices let us test the Central Dogma’s information flow.

We fitted multiple linear regression models, where the focal dependent variable could be gbM, mRNA or protein, across various subsets of expressed genes, and the independent variables the lower omic levels measured in the same genes. For this modelling we reparametrized CDS codon frequencies as the combination of three, biologically interpretable, nested components of increasing complexity, namely protein length (the sum of all codon frequencies), then the 20 amino acid frequencies (the sums of frequencies for those codons representing a given amino acid, requiring 19 additional parameters), and finally codon usage within each amino acid (representing the deviations from the baseline effect for an amino acid, comprising 41 extra parameters). The first two of these components are linear combinations of the codon frequency counts, and the third accounts for variation due to the choice of codon within an amino acid class. For clarity, we refer to “codon frequencies” when interpreting CDS as a model of the 61 codon counts (i.e. as in the previous sections), and “codon usage” when interpreting it after separating out protein length and amino acid frequency. Thus, depending on the parameterisation used, the same fitted model can be reinterpreted to provide different insights, while yielding identical predicted effects and explaining the same total variance. This reparameterization was used to test the effects of adding increasing information about sequence composition on constitutive gbM, mRNA or protein expression within a single analysis of variance.

Each model is expressed in the form *Y ∼ A+B+….*, where *Y* is the target omic level and *A, B* … are explanatory omic levels. For example, the model *Col-protein ∼ CDS + gbM + mRNA*means that protein expression across the genes with protein and mRNA expression in Col-0 is modelled in terms of first codon frequencies (CDS), then gene body methylation (gbM) and finally mRNA expression measured in those genes. The order in which levels are included in a model affects how much variation is explained by each level (i.e. the modelling is greedy, so each level is assigned the maximum possible variation after allowing for the previously fitted levels) thereby revealing confounding between levels. For example, fitting gbM either alone or after fitting CDS reveals how much variation in gene expression is solely attributable to gbM. We also subdivided the genes into two classes; those 7,771 with both measurable mRNA and protein expression; – the difference is due to a few genes with multiple annotated stop codons which were excluded), and those 9,633 with only mRNA expression and denoted mRNA* or gbM* in the **Figure 9** and **Supplementary Table S8**. Both mRNA and protein expression were log-transformed prior to fitting the linear models which therefore represent multiplicative effects on expression. The **Figure 9** legend describes the dependent and explanatory omic levels in detail.

**Figure 9.**
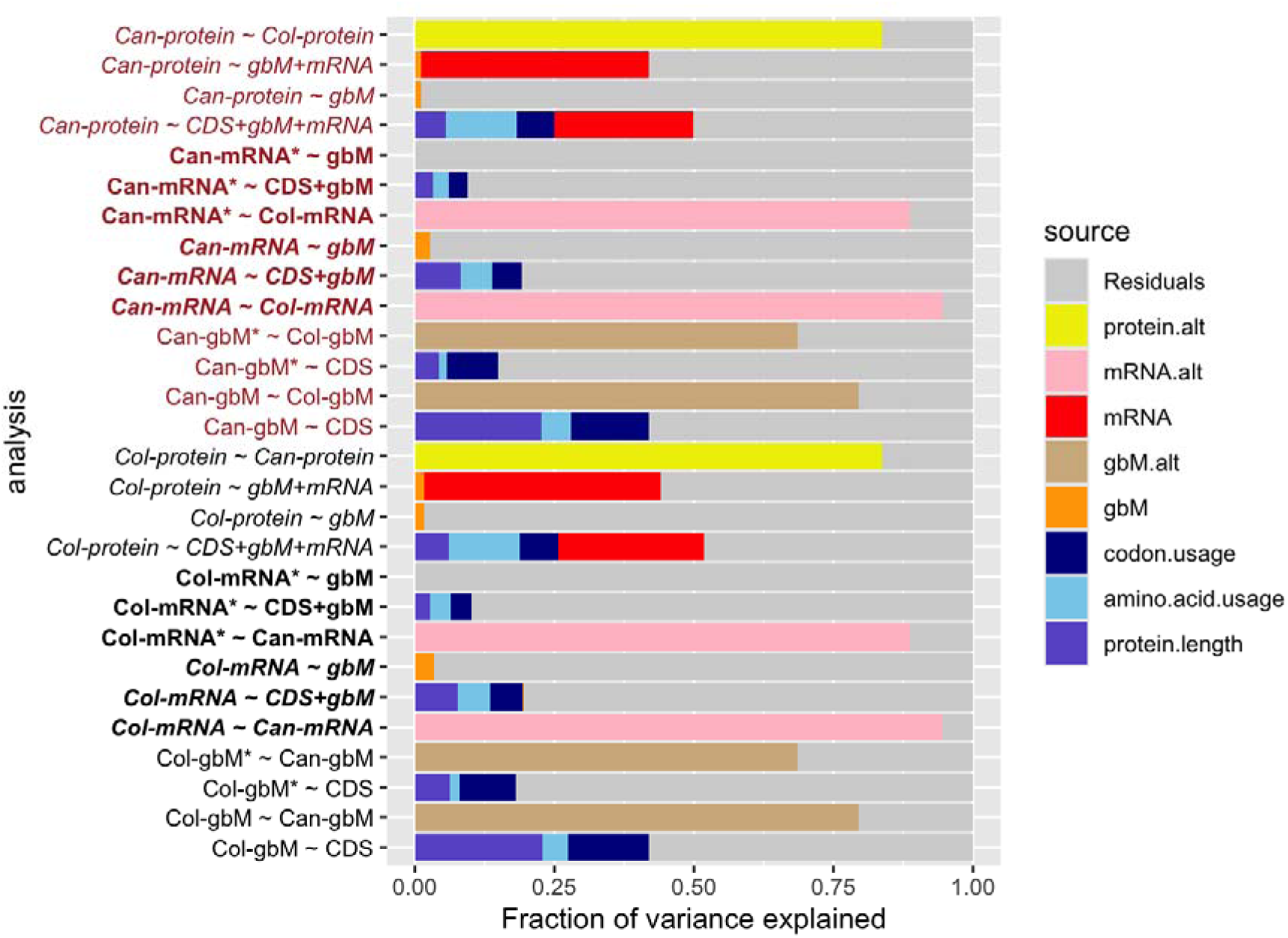
Bar plots of variance explained by multiple linear regression models. Each row represents one model. The model is specified on the left, the colour indicating whether Col-0 (black) or Can-0 (brown) is the target omic level (dependent variable). Each model is by the formula Y ∼ A+B+…, where Y is the target and A, B … are explanatory omic levels. The targets for the Col-0 analyses are:- Col-protein: log-transformed protein expression, Col-mRNA: log-transformed mRNA expression for genes also with protein expression; Col-mRNA*: log-transformed mRNA expression for genes without protein expression; Col-gbM: percent gene-body methylation for genes also with protein expression; Col-gbM*: percent gene-body methylation for genes without protein expression. Similar names apply for the Can-0 analyses. The explanatory omic levels are:- CDS: coding DNA sequence composition, (partitioned into protein length, amino-acid usage and codon usage); gbM: percent gene body methylation; mRNA: log-transformed mRNA expression; gbM.alt, mRNA.alt, protein.alt: expression of gbM/mRNA/protein in the alternative accession (i.e. Can-0 if the target accession is Col-0). The barplot for each analysis represented the fraction of variance explained by each term in the model, using the colour-coding given in the legend. CDS effects are partitioned into protein.length, amino.acid.usage and codon.usage; The horizontal extent of each bar represents the fraction of variance due to the corresponding variable, after first fitting the preceding variable sin the formula from left to right.

The analyses are summarised as barplots in **Figure 9**, where the horizontal extent of each bar indicates the fraction of variance in the target omic level attributable to the corresponding component in the model, after fitting the preceding terms. For comparison we also show the results of modelling an omic level in one accession by the corresponding level in the alternate accession (pink bars: mRNA and yellow bars: protein). The results for Col-0 (**Figure 9** upper) and Can-0 (**Figure 9** lower) are extremely similar, illustrating the robustness of these analyses to genetic perturbation.

The p-value of each variance component from its corresponding partial F-test is given in **Supplementary Table S8;** virtually all components are extremely significant with analysis of variance p-values often much smaller than 10 even when the fraction of variance explained is too small to be visible. That is, statistical significance is necessary but not sufficient to imply biological importance. Multiple linear regression models we use here are “greedy”: the order in which explanatory variables are fitted in the model determines how much variance each explains. Comparing models in which the same variables are added in different orders reveals statistical confounding, i.e. when the same outcome is attributable to different causes.

We consider the impact of explanatory omic levels on expression in Central Dogma order. The simplest explanatory variable, protein length, is known to be anticorrelated with expression [35]. In Col-0, we find protein length alone explains 7.7% of the variation in mRNA expression, (double that explained by gbM at 3.5%), and 6% of protein expression variation. The statistics in Can-0 are similar (mRNA: 8.1%, protein: 11.1%). The next component of sequence composition, amino-acid usage, explains significant additional variance in mRNA (5.7%, 5.8%) and in protein expression (12.8%, 12.7%), after accounting for sequence length, although spread across 19 parameters. Codon usage explains further variance (mRNA:6.0%, 5.3%; protein: 6.8%,6.7%), but spread over far more (41) estimated parameters. Overall, CDS effects on mRNA expression (19.5%, 19.2%), are lower than protein expression (25.6%, 24.9%), and the relative impacts of the three CDS components also differ. When we model the mRNA expression of those genes without protein expression (denoted mRNA* in **Figure 9**) the fractions of variance explained by CDS are halved (10.1%, 9.4%).

We then modelled constitutive gbM as a function of CDS. Among genes with both mRNA and protein expression we find CDS effects account for close to half (42.0%, 42.0%) of gbM variance. However, these fractions are more than halved (18.1%, 14.8%) among genes with only mRNA expression. Modelling gbM in one accession by the corresponding level in the alternative accession (shown as tan coloured bars in **Figure 9**) explains far more of the variance (79.4% for genes with protein and 68.6% for those without) and shows that constitutive gbM is highly reproducible (as expected, given the reproducibility of the underlying CpG methylations reported above) but that some of this reproducible variation is unexplained by codon frequencies.

We next treated gbM as an explanatory level to model mRNA and protein expression. When considered in isolation (i.e., excluding CDS effects), gbM has a relatively small but highly significant impact on mRNA expression in genes with both protein and mRNA expression, explaining (3.5%, 4.2%) of mRNA variance, but a negligible impact on the expression of genes without protein (0.1%, 0.1%). When CDS effects are fitted before considering gbM, virtually all the effects of gbM are ablated; under a constant environment, the impact of constitutive gbM on mRNA expression mediates a small fraction of sequence composition effects.

When modelling protein expression, if mRNA expression is added after CDS and gbM, in total half of the variation in protein expression can be explained (51.9%, 49.8%), and which exceeds that when only gbM and mRNA are included (44.1%, 42.0%). Thus, part of the information encoded in CDS relevant to protein expression is not mediated through mRNA, in contradiction to the Central Dogma. Similarly to modelling gbM, much greater fractions of variation are explained by modelling Col-0 mRNA by Can-0 mRNA (and vice versa) (94.5%, 94.5%) or Col-0 protein by Can-0 protein (83.8%, 83.8%). Thus, whilst our simple regression models are powerful, they do not capture all the information, and there is unexplained yet reproducible variation.

### Differential comparisons of omic levels between accessions

We observed 7,585 differentially expressed (DE) mRNAs at FDR < 0.05 among the 17,771 orthologous Col-0:Can-0 gene pairs at which we could make a determination, ignoring protein expression status. In the subset of 7,060 genes with differential determinations for both mRNA and protein, we observed 866 DE proteins (FDR<0.05) and 2,850 DE mRNAs (FDR<0.05; EdgeR did not determine DE status for all genes, so these subsets are slightly smaller than those in the previous sections) (**Supplemental Table S9**). **Figure 10 A** plots the log_2_ fold change in expression (logFC) for mRNA vs protein, color-coded by DE FDR. It shows that logFC is broadly consistent between mRNA and protein, although there are a significant number of genes which are DE for only mRNA or only protein. Of the DE mRNAs and DE proteins, 579 (20% of DE mRNAs and 67% of DE proteins) are in common (Fisher’s Exact test < 10^-32^), as shown in the Venn Diagram in **Figure 10 B**. Gene Ontology (GO) enrichment analysis [36] revealed distinct but overlapping enrichments between DEGs and DEPs (**Supplemental Figure S7**). Proteins and transcripts associated with glucosinolate biosynthesis exhibited increased abundance in Can-0, whereas proteins associated with immunity (“hypersensitive response”, “cell death”, “response to biotic stimulus”) and abscission were increased in abundance in Col-0 (**Supplemental Fig. 7**.

**Figure 10.**
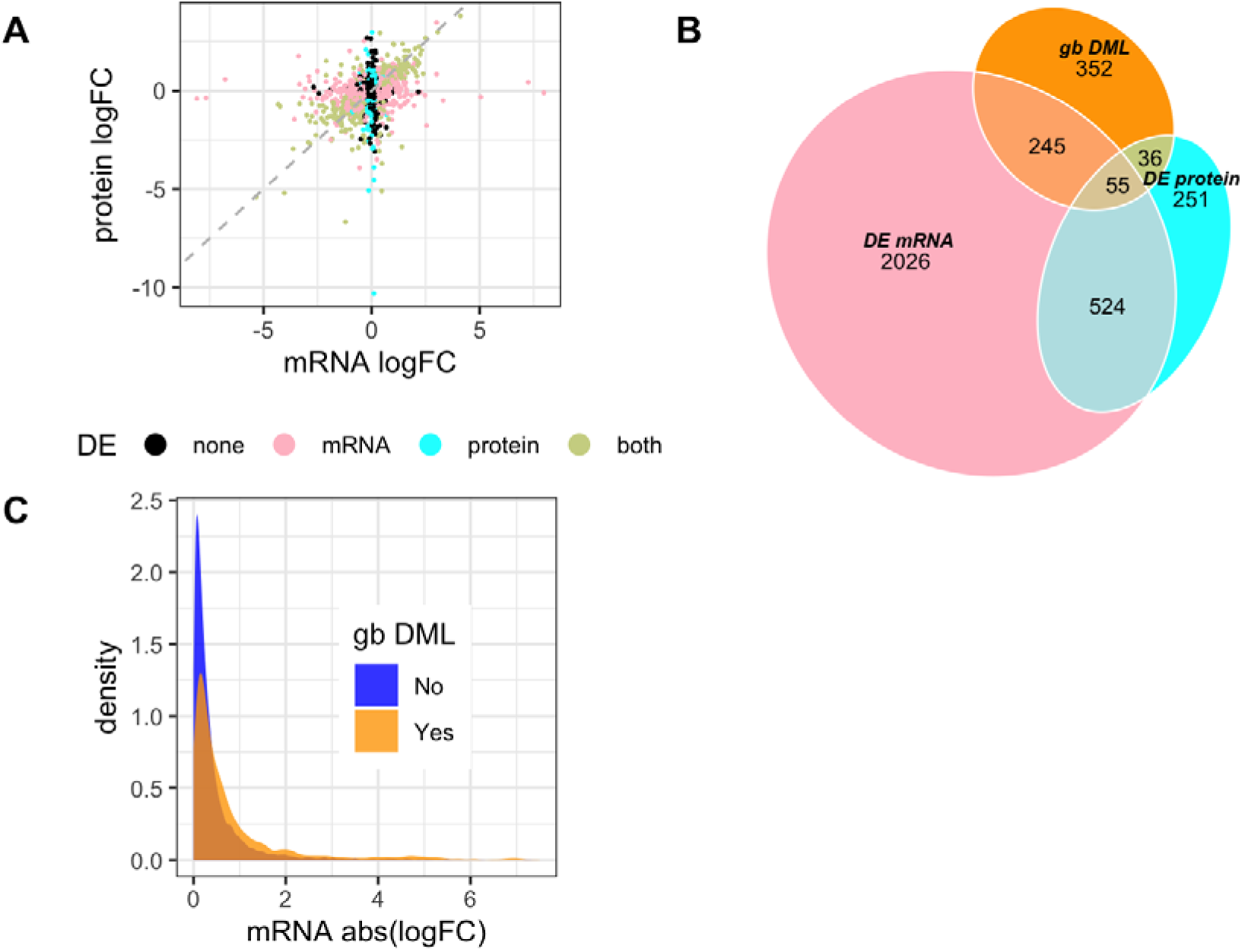
Differential expression (DE) and differential gene-body methylation (gb DML). (A): Scatter plots of log_2_ fold change (logFC) for mRNA (x-axis) vs protein (y-axis) for 5,698 ortholog pairs between Col-0 and Can-0 with DE determinations in protein and mRNA. Points are color-coded according to whether the pairs are DE at both protein and mRNA, only protein, only mRNA or neither (all determinations using FDR<0.05). (B): Venn diagram of overlaps of mRNA and protein DE and gb DML. Numbers are counts of DE gene pairs within each subset (e.g. there are 2,026 pairs that are only DE mRNA, 524+55 = 579 that are both DE mRNA and protein, and 55 that are DE mRNA and protein and gb DML). (C) Distributions of absolute log2 fold-change in mRNA expression between Col-0 and Can-0, for genes with (orange) or without (blue) gb DML.

### Genetic and epigenetic correlates of differential expression

We next characterised how differential mRNA or protein expression related to differences in CDS and to CpG methylation, either outside or inside the gene body. We distinguished between a differentially methylated locus, DML - a single syntenic CpG dinucleotide at which methylation differs between Col-0 and Can-0 - and a differentially methylated region (DMR), in which average methylation differs across the CpG dinucleotides within the region. If a gene body overlaps with at least one DML or DMR, this gene body is defined as with DML or DMR. We used the same definition to classify intron and exon regions, and genomic contexts up- or down-stream of the gene body, and for structural variants (SV) as discussed below. The Venn diagram in Figure 10B shows the overlaps with gb DML genes at 5% FDR. Of the 688 genes with gb DML, 336 (49%) are DE for mRNA or protein, or both.

We compared DML and DMR gene classifications with the corresponding absolute values of logFC expression, reporting −log_10_ p-values (logP) of the Mann-Whitney tests, which are robust non-parametric tests of differences in the average ranks of the absolute logFC of expression between genes with or without differential methylation. This analysis therefore does not require differential transcript or protein expression at any specific FDR threshold but instead considers trends. The results are summarised in **Figure 11 (Supplemental Table S9).**

**Figure 11.**
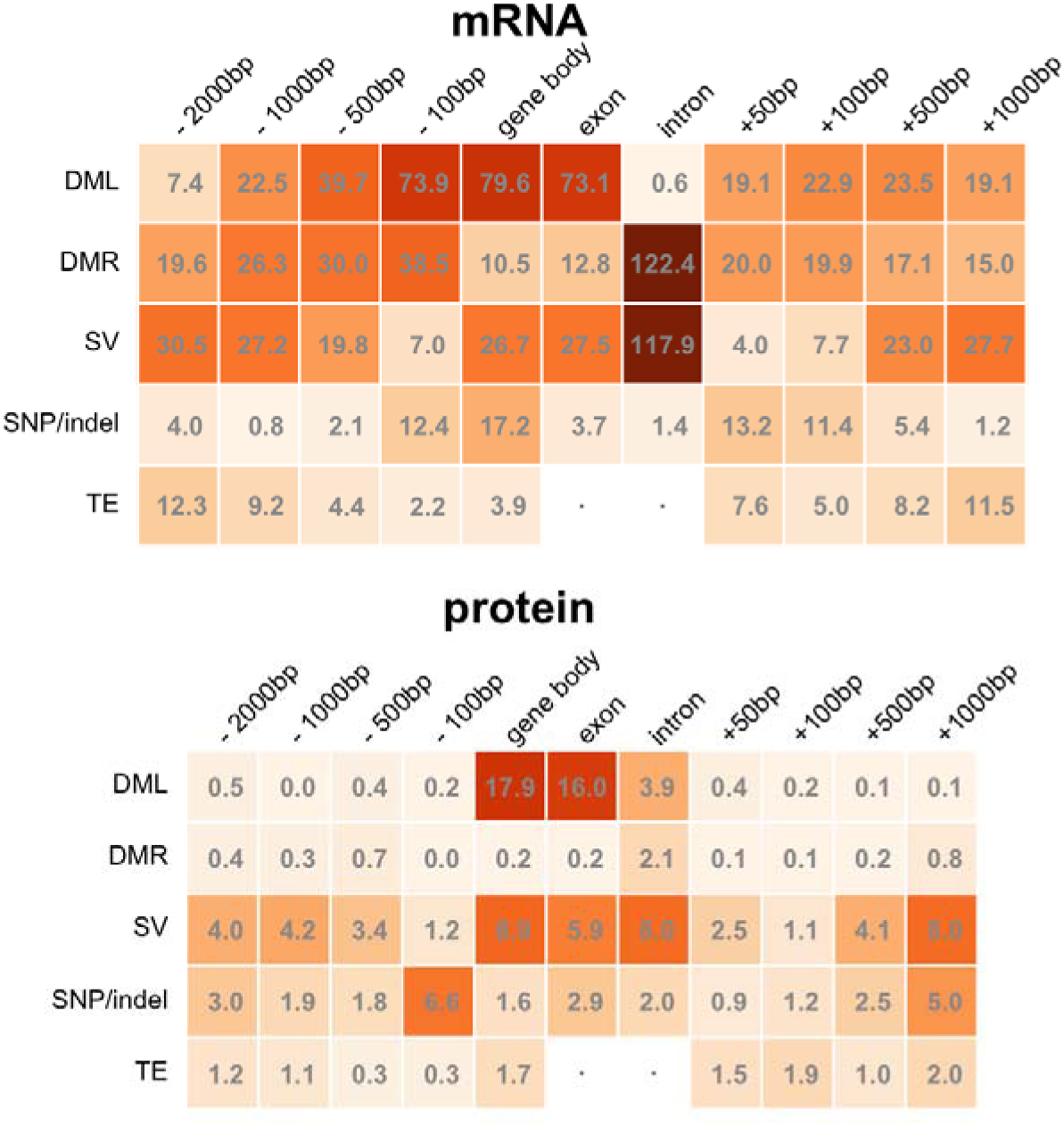
Impact of differential methylation, structural variation, and indels on differential (A) mRNA and (B) protein expression between Col-0 and Can-0. The x-axis represents a schematic gene, comprising upstream, gene-body (subdivided into intronic and exonic components) and downstream genomic contexts. The y-axis represents the variation categories DML: differentially methylated loci, DMR: differentially methylated regions, SV: structural variations, indels: short insertion-deletions, TE: transposable elements. The number in each is the negative log10 p-value of the unsigned Mann-Whitney test of association between the category in the given genomic context and differential expression, (except for TE which shows the logP of the Spearman rank correlation test reflecting the fact that differential TE abundance is measured quantitatively). The orange shade of the cell indicates the strength of association, from dark (strong) to pale (weak).

Despite the high correlation of gbM between Col-0 and Can-0 at orthologous genes (), the presence of DML or DMR within or nearby a gene body strongly associates with absolute log-fold changes in mRNA expression (**Figure 11).** DML and DMR in gene bodies, or within 100 bp upstream, have the highest impact on differential mRNA expression. In general, the presence of even a single locus methylation difference (i.e. DML) is a stronger predictor of gene expression difference than is DMR, with the exception of intronic DML which is not significant for mRNA abundance (logP = 0.61), but highly significant for intronic DMR (logP = 122.42). **Figure 10B** show the overlaps between DE mRNAs and proteins and gb DML. We found that the corresponding signed Mann-Whitney tests (ie where we did not take absolute values of logFC) were markedly less significant. **Figure 10C** shows the distributions of absolute log-fold changes for mRNA expression for genes with or without gb DML. Although the distributions appear broadly similar, they have highly a significantly different Mann-Whitney statistic (logP=79). Thus, although differential methylation is strongly statistically associated with differential expression, it is not a reliable predictor.

We then used the same methodology to test if sequence differences are associated with differential mRNA and protein expression. We tested for associations between the presence of SNPs or small (< 10 bp) indels, or structural variations (SV, defined as indels>10bp) nearby or within differentially expressed genes as for methylation using Mann-Whitney tests. Gene-body SVs, especially in introns, have strong associations on differential mRNA abundance (Figure 11A). Interestingly, distant structural variants can have stronger associations with differential gene expression. Similar if less significant patterns occur for small variants.

## Discussion

It is a remarkable fact that although most genes are encoded only once in the nuclear genome, their constitutive expression levels in a given tissue vary by orders of magnitude [22]. In the *Arabidopsis thaliana* rosettes studied here, the most highly expressed genes exceed the median level by over 1,000-fold for mRNA and over 500-fold for protein. An important question is how these levels are set and maintained, and their reproducibility. We have shown here that in two genetically divergent accessions of *Arabidopsis thaliana,*these levels are indeed highly reproducible between biological replicates in a controlled environment, and that a simple multiple linear regression model based on gene codon frequencies is unexpectedly powerful at modelling both mRNA and protein expression levels. A gene’s constitutive expression is thus partially determined by its internal codon composition, which presumably evolved to express it at its optimal level by selecting codons adaptively and tuning tRNA abundance to match overall demand for amino acids. The diagram in **Figure 12** summarises our findings.

**Figure 12.**
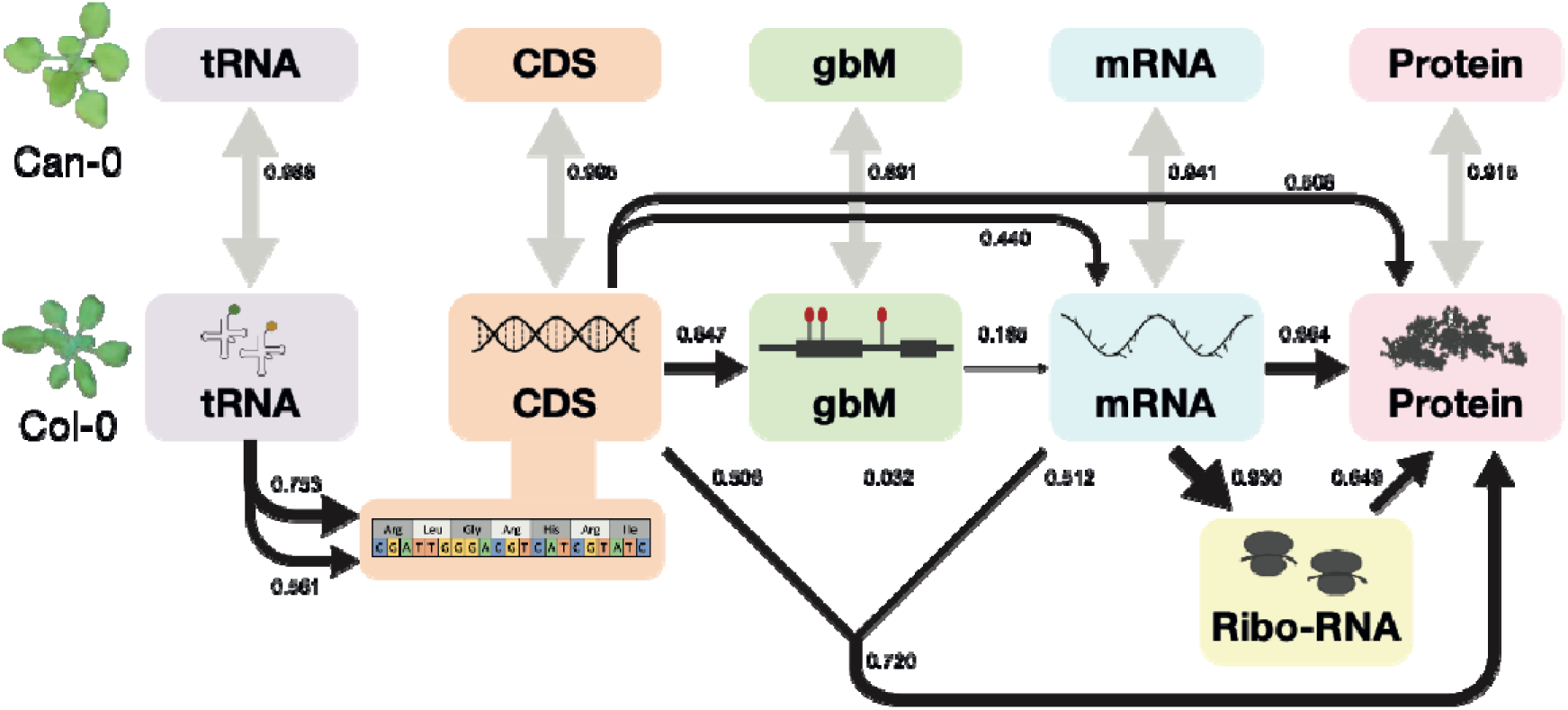
Genomic information pathways and their numerical linkages as observed in this study, in relation to the Central Dogma. The boxes show different omic levels in Col-0 and Can-0. Black arrows indicate the flow of information implied by the Central Dogma starting from CDS (peach) via gbM (green) to mRNA (light blue), ribosome-associated mRNA (yellow) and finally protein (pink). The strength of Pearson correlation between levels is shown both by the numbers and by the thickness of the corresponding black arrows, and represents the correlation between quantitative expression levels (except in the case of CDS where is represents the correlation derived from the multiple regression codon frequency model, i.e. the square root of the fraction of variance explained). The merged arrows connecting CDS, gbM and mRNA to protein show the result of combining information into a single model; note that their combined correlation of 0.720 is less than the sum of their individual effects. The correlation of tRNA levels (lilac) with genome-wide codon and amino acid frequencies is shown on the left. The correlations between corresponding Col-0 and Can-0 omic levels are shown next to the grey double-headed arrows.

Our results are based on a comparison of just two accessions in a single tissue, and under controlled environment. They should be extended to a wider set of genomes and across different tissues and environments to take account of expression quantitative trait loci, cell-type effects and environmental variation. The two genomes used here are from the set of 19 founders of the Arabidopsis MAGIC population of recombinant inbred lines descended from these founders [14, 37]. Work is underway by our group to analyse omic levels across the founders and the MAGIC population, which will enable us to test if our conclusions are robust in the presence of significantly more genetic variation.

### Codon and tRNA Effects on Expression

Codon frequencies alone account for about 19% of the variance of mRNA expression, and about 25% of protein expression, among those *7,771* genes with measurable mRNA and protein expression in both accessions. Augmenting the codon model of protein expression with mRNA expression data almost doubles the total to about 46%. Interestingly codon frequencies only explain 9% of variance among the 9,633 genes with mRNA but no protein expression, suggesting the expression of these genes is controlled by different factors.

When interpreting these results, it helps to bear in mind that the total fraction of variance explained by a model equals the squared correlation between the observed and fitted values. That is, correlations are larger than their equivalent variance fractions; 19% variance is equivalent to a correlation of *0.46*. In **Figure 12** all the numerical linkages between omic levels are shown on the correlation scale. However, it is more meaningful to report variance fractions when decomposing a multi-component model in an analysis of variance (**Figure 9**). In addition, we report results after log-transforming expression, so our models are multiplicative on the original measurement scale. This means the impact on the original scale of expression of the codon *c* with gene frequency *n*, will be proportional to 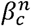, where *β_c_* is both the regression coefficient we estimate and the multiplicative impact of a single copy of the codon on the original measurement scale.

If full sequence information from each gene is used for prediction (i.e. including the order of bases), instead of being summarised as the 61 codon frequencies as was done here, it is possible to train large language models with millions of parameters to predict expression patterns. In a different Arabidopsis data set, and encoding each gene sequence by its sequence of codons, mRNA prediction accuracies of between *R*^2^ = 0.2 - 0.4 were achieved, depending on the model used [7]. Thus, there is additional, exploitable, information encoded in the order of the codons in each gene. Nonetheless it is remarkable how well our simple, biologically interpretable, model performs.

We find that a given codon has similar impacts on mRNA and protein expression; the correlations between the 61 non-terminator codon effects on mRNA vs protein expression (*R* = 0.80, **Figure 6 A,B**) exceed those of the actual expression measures (*R* = 0.64 - 0.66, **Figure 4 A,B**), suggesting that an underlying biological mechanism is being isolated. Whilst codon effects could point towards tRNAs as mediators for their effects on protein translation, our analyses are also consistent with the hypothesis that codons which increase protein translation also reduce mRNA decay, driving mRNA stability and hence mRNA abundance [38–40]. Interestingly, we find that tRNA abundance is uncorrelated with these estimated codon effects on expression but instead correlates with overall codon abundance (**Figure 8 B,C**). Furthermore, the aggregate tRNA abundance for all tRNAs specifying a given amino acid (ie isoacceptors) tracks the overall frequency of that amino acid across the proteome even more strongly than at the codon level (**Figure 8 D**). Our results suggest that tRNA abundance adapts dynamically to the overall demand for amino acids but that differences in translation efficiencies between tRNAs do not measurably affect abundance of specific genes. The correlations we observed between tRNA abundance and codon usage resemble those reported in humans [31]. In summary, codons interact with two distinct phenomena – their abundance relates to tRNA abundance, but independently of their impact on expression.

Our results are broadly consistent with studies in human cell lines [41], where a combination of sequence features (including protein length and predicted mRNA decay rate) and mRNA expression explained about two thirds of the variance in expression of 512 proteins. Another study of mRNA half-lives in human and mouse [42] also identified codon frequencies and protein length as key factors. Although we did not measure mRNA decay rates, it is likely that our models of mRNA levels are implicitly modelling them. In fact, of the 18 codons listed in [37] as impacting mRNA half-lives in humans, our data in Supplemental Figure S4 share the same sign in at least 16 cases, a statistically significant coincidence (*P* < 0.008, See **Supplemental Data File S7)**. Our data were all collected under uniform unstressed conditions, so we could not measure the impact of stress on codon usage, which affects the translation of specific codons [43].

We exploited the fact that the codon frequency model is mathematically equivalent to the combination of three simpler models based on protein length, amino-acid frequency, and codon usage. We find mRNA expression decreases with gene length, and the choice of encoded amino acids and the choice of codon each have significant but smaller impacts than protein length. The relative impacts of these factors on protein expression are subtly different: for example, gene length is less important than amino acid choice. In addition, among genes with mRNA but no protein expression the impacts of all three factors are much reduced (**Figure 9**).

Interestingly, a codon’s impact on expression is entirely unrelated to its genome wide frequency among expressed genes; it is not true that highly expressed genes preferentially use high frequency codons. Therefore, to increase the expression of a protein by editing its codon composition, in general one should select optimal codons based on their effects estimated from a model like that fitted here, but derived from expression data from the species of interest. If the close similarity of codon effects on mRNA and protein expression that we observed generalises to other species, then it would be sufficient to train codon models on mRNA expression alone, which is experimentally more tractable. However, it is important to note that whilst the codon models explain a significant proportion of mRNA and protein expression, there will be important nuanced effects of codon usage for individual genes. Synonymous substitutions can influence diverse mechanisms related to gene expression and protein homeostasis including transcriptional regulation, mRNA lifetime, translation initiation efficiency, translation elongation rate and downstream effects on protein folding as well as degradation [27].

### Constitutive Gene Body Methylation Effects

Our results minimise the role of constitutive gene-body methylation in regulating expression[12]. We found that, notwithstanding the high degree of conservation of gbM observed between accessions, when the environment is controlled, augmenting the codon models of mRNA or protein expression with constitutive gbM data makes only a negligible improvement to model fit. Indeed, constitutive gbM itself is largely predicted by local codon frequencies, explaining about 44% of gbM variation. Under these environmental conditions, constitutive gbM potentially mediates some of the impact of codon frequencies – explaining about 3-4% of mRNA and protein variation in the absence of codon frequency data - but does not contribute new information over that encoded in CDS. Since constitutive gbM is highly conserved between accessions it is indeed plausible that it is driven by local sequence context.

Interestingly, many codons with strong effects on gbM do not contain CpG dinucleotides. This suggests that non-CpG local sequence context drives CpG gbM. Additionally, whilst protein length is the major determinant of gbM variation in genes that exhibit both protein and mRNA expression, it is irrelevant for genes with only mRNA expression, illustrating major differences in the genetic architecture of gbM depending on protein expression.

However, we caution against the view that constitutive gbM is irrelevant to expression for the following reasons. First, among genes with mRNA expression in rosettes, the distribution of gbM is concentrated around 10% (**Figure 5 A,B**) with little variation and hence limited opportunity to influence expression. However, there is a distinct subset of several hundred centromere-associated genes without any mRNA or protein expression in rosettes for which gbM exceeds 75%. The expression of these genes might be actively silenced by these high gbM levels, but this does not explain how a further ∼5,000 genes are neither expressed nor highly methylated, and indeed these highly methylated genes may simply be passively reflecting local methylation levels around the centromeres. Second, differential methylation, both in gene bodies and elsewhere, is strongly associated with differential mRNA expression between Col-0 and Can-0. Interestingly, the presence of a single differentially methylated CpG (DML) is a better indicator of DE than is the average difference of methylation (DMR), and moreover the direction of the change in expression is uncertain; unsigned association tests which ignore the direction of the DE are more significant than signed tests. DML indicate perturbations in methylation due to local sequence differences, which increase the variance in expression rather than its direction **(Figure 10 C**). Differential methylation outside of gene bodies affects mRNA expression less than does gbM and has markedly less impact on protein expression (**Figure 11**). Differences in transposable elements between the accessions are only weakly associated with differential expression of nearby genes (**Figure 11)**.

### Genes that express protein and mRNA differ from those that only express mRNA

The distribution of mRNA expression is very different for genes which also have protein expression, compared to those for which protein was not detected in our proteomics workflow. It is not true that moderately high mRNA expression necessarily implies any protein expression (**Figure 4**). In fact, many genes without protein expression have higher mRNA levels than those with protein. This discordance has been noted in numerous organisms [1–6]. Those genes with both mRNA and protein have an overall distribution close to lognormal. Taken together with the log-transformations used here, this suggests that these genes are being expressed as the result of a balance between independent multiplicative stochastic processes of synthesis and decay. In contrast, genes without protein expression have a more complex distribution.

Surprisingly, using ribosome-associated mRNA expression levels does not change the picture, at least in regard to genes with both protein and mRNA expression. The only noteworthy difference appears in genes with mRNA expression but not protein – about 30% of which do not appear to be associated with the ribosome. However, it should be noted that 3’RiboSeq merely quantifies transcripts associated with ribosomes, without distinguishing between monosomes and polysomes and does not indicate whether the transcripts are actively translated. It may be that stronger correlations with protein expression translatome data could be obtained from ribosome profiling in which it is possible to accurately determine the average number of ribosomes per mRNA and thus estimate the relative translation levels of a transcript [44].

We observed more genes with differential mRNA than protein expression between accessions. To some extent, this may reflect the limitations of quantitative proteomics which do not readily permit sampling of the entire proteome, but it may also indicate buffering of the proteome. The discrepancy may also be related to the fact that protein expression is estimated (here) from peptide DIA data and is therefore subject to different measurement issues and different algorithmic processing steps than RNAseq data.. If this step introduces biases, they are likely to be the same within genes across replicates, thereby contributing to differences between genes, and consequently mRNA/protein correlations. Even with these experimental and methodological issues, it is remarkable how estimated codon mRNA expression effects resemble protein effects (**Figure 6**).

### The best predictor of expression in one accession is expression in a different accession

Despite the success of the codon models in **Figure 9**, the expression of gbM, mRNA or protein in a given focal accession is more strongly correlated with orthologous expression in the other, genetically distant, accession than with lower omic levels within the focal accession. This orthologous expression fidelity remains unexplained. Possibly this is due to active homeostatic feedback mechanisms and the conserved effects of transcription factors on the control of expression under unstressed conditions. In a future study, it will be interesting to determine to what extent this predictive power is maintained in the presence of a stimulus such as a stress or a developmental cue. Another interesting question is what underlies the residual variation in constitutive protein and mRNA expression that is unexplained by the models used here. This is not due to measurement noise, because the reproducibility between biological replicates is so high. Rather, this might reflect factors such as differences in protein and mRNA degradation, related to the adaptation of the two accessions to different environments.

### Long-read genome assemblies and annotations are accurate but not yet perfect

Comparisons of our assemblies of Col-0 and Can-0 with published long-read assemblies of these accessions show that although they agree with high fidelity over most of the genome, some differences, concentrated within tandem repetitive regions remain. Typically, assemblies of the reference accession Col-0 disagree at about a few thousand SNP positions (**Figure 3**). These discrepancies are partially due to algorithmic differences in the assembly pipelines but might also reflect some genuine variation in the germplasm sequenced in each study. Similarly, the identification of gene orthologs between accessions depends on how paralogy within an accession is defined; orthologous genes can have different numbers of isoforms and so are arguably no longer true orthologs.

### The Central Dogma Revisited

Our study is incompatible with a naive interpretation of the Central Dogma’s flow of information; it is not true that all the relevant information in CDS passes though mRNA abundance levels alone to modulate protein levels. One potential explanation is that our measurements of mRNA and protein expression are at single time-points and represent the difference between synthesis and degradation integrated over the recent past. Potentially more information would be available if expression were measured at different time points due to e.g. diurnal cycles. Some alternative explanations seem unlikely: First, our ribo-Seq levels are no better correlated with protein abundance than are standard RNA-seq levels, providing no support for the hypothesis that we are somehow biased away from relevant mRNA expression measurements by including transcripts not associated with ribosomes. Second, our data are from bulk tissue and not single cells. If it were technically possible to measure mRNA and protein from the same cells we might observe stronger correlation between mRNA and protein, but it is difficult to see how this would explain how the additional information in CDS is actioned, because the latter is constant across cells. Third, tRNA abundance data cannot explain the discrepancy.

Our CDS modelling summarises just the gene codon frequencies, implicitly modelling gene length as well as gene amino acid frequencies and codon usage bias. As **Figure 9** shows, all three of these components are informative. We must therefore conclude that these codon frequencies are relevant to protein expression but in a manner not mediated by transcript levels alone.

It is known that on average protein and mRNA expression reduces with gene length [32], and our data supports this, but like other studies we report mRNA and protein abundance levels as normalised estimates of the numbers of copies of these molecules, independent of their length. This might explain why the component of CDS due to gene length is so relevant in our models. We do not have a complete mechanistic explanation for why amino-acid frequencies and codon usage are also important. Regardless, since all the codon information in a CDS is also present in the sequence of its mRNA transcript, the Central Dogma still holds if the definition of information includes both abundance level and sequence. In a sense this is closer to Frances Crick’s original conception [45].

### Conclusions

This study demonstrates how simple sequence features underlie much of the variation in expression of different omic levels and, in part, how these levels depend on each other, when and where it is helpful to measure a level, and when we can substitute one that is difficult to measure by a more tractable alternative. In particular, we have shown that, in the absence of environmental stress, the levels of constitutive methylation and tRNA abundance and their effects on expression are consequences of underlying sequence features. Finally, the Central Dogma’s flow of information must be treated as multidimensional.

## Declarations

**All manuscripts must contain the following sections under the heading ‘Declarations’:**

## Ethics approval and consent to participate

Not applicable

## Consent for publication

Not applicable

## Availability of data and materials

All Col-0 and Can-0 DNA and RNA sequence data and the genome assemblies are publicly available from ENA under the study accession number ERP161680. All samples and their accession numbers are listed in **Supplementary Table S10**.

The mass spectrometry proteomics data have been deposited to the ProteomeXchange Consortium via the PRIDE [46] partner repository with the dataset identifier PXD058342.

## Reviewer access details

Log in to the PRIDE website using the following details:

## Project accession: PXD058342 Token: NaaHovzhdBRn

Alternatively, reviewers can access the dataset by logging in to the PRIDE website using the following account details: Username: Password: iHQH1QnAEHxj

## Supplementary files

R code that generates the Figures

## Authors’ contributions

**FLT, KSL, RM, KHP, GS, DN** conceived the project and designed the experiments

**MB, YK, NPA, BP, CR** performed the experiments

**ZZ, RM, YK, LA, KHP, DB** performed the analyses

**RM, FLT, KSL, ZZ, YK, XL** wrote or edited the paper

## Acknowledgements

We thank Hongtao Zhang for preliminary data to support the funding application and Rob King for preliminary RNA-seq analysis. We also thank the Earlham Institute and the UCL Long Read Sequencing Facility for DNA and RNA sequencing and are grateful to Ian Henderson and Oxford Nanopore Technology technical support staff for advice on long read sequencing and genome assembly.

## Methods

### Plant growth

Seeds of the *Arabidopsis thaliana* accessions Col-0 and Can-0 were obtained from the Eurasian Stock Centre (formerly NASC). Seeds were sown on pre-soaked Levington F2 plus sand mix and stratified for 5 d at 5 °C before transferring to a controlled environment room. For long read whole genome sequencing, plants were grown in short-day conditions: 10 h under fluorescent bulbs at 250 µmol/m^2^ /s, 23°C and 65 % relative humidity, followed by 14 h darkness at 18°C with 75% relative humidity, and harvested shortly before bolting. Unless otherwise indicated, for all other assays, plants were grown in long day conditions: 16 h under LED lights at 150 µmol/m^2^ /s, 22°C and 65% relative humidity, followed by 8 h darkness at 18°C with 75% relative humidity and harvested at the 9-leaf stage.

### Illumina whole genome DNA sequencing

DNA from a single leaf of each accession was extracted using the DNeasy Plant Mini kit (Qiagen, Manchester, UK) with on-column RNase treatment. Library preparation and Illumina sequencing (150 bp paired end reads) was performed by the Earlham Institute, UK. Read quality was checked using *Fastqc* [47], and sequencing adapters were trimmed using *BBduk* [48] with options ‘*ktrim=r k=23 mink=11 hdist=1 tpe tbo*’. To retain only good quality reads, a further round of quality trimming was conducted by *BBduk* with options ‘*qtrim=r trimq=10 minlen=50*’.

### Long-read whole genome DNA sequence

Col-0 and Can-0 plants were grown under short day conditions until shortly before bolting and a single rosette for each genotype (∼2 g tissue) was harvested and snap frozen in liquid nitrogen. High-molecular weight (HMW) DNA was extracted using a NucleoBond HMW DNA extraction kit (Macherey-Nagel, Dueren, Germany) as per the manufacturer’s instructions. DNA quality control was performed using agarose gel electrophoresis, UV spectrophotometry (NanoDrop Technologies, USA), and the FP-1002 Genomic DNA 165 kb kit for FEMTO Pulse systems (Agilent technologies, Stockport, UK). DNA was quantified using Qubit high sensitivity DNA quantification kit (Thermo Fisher, Altrincham, UK). HMW DNA was sent to the Long Read Sequencing Facility (LRS) at University College London (London, UK) for library preparation and sequencing.

Libraries for sequencing using Oxford Nanopore Technologies (Oxford, UK, hereafter abbreviated to ONT or Nanopore) were prepared by using the ONT SQK-LSK109 ligation kit and sequenced on an ONT PromethION instrument. Initial sequencing of Can-0 produced an excessive number of short reads (< 10 kbp), therefore the Short Read Eliminator Kit (SS-100-101-01; Circulomics Inc, USA) was used to progressively remove reads <25 kbp in the Col-0 library before sequencing. Basecalling for ONT data was conducted using *guppy_basecaller*function in *Guppy* 5.0.16 [49], with options ‘*-c dna_r9.4.1_450bps_sup_prom.cfg --min_qscore 9 --min_score 40 --trim_barcodes*’. We obtained 22.5 Gb data for Can-0 (150x), with a read N50 value around 26 kbp, and 30.3 Gb data (202x) for Col-0, with read N50 value around 33 kbp.

LRS also prepared libraries for PacBio HiFi sequencing (Pacific Biosciences of California, Inc. hereafter PacBio HiFi). DNA fragments were firstly sheared to ∼17 kb by using Megaruptor 3 (Diagenode Inc.), and then the SMRTbell® Express Template Prep Kit 2.0 was used for constructing sequencing libraries. Libraries were sequenced on a PacBio Sequel II. Basecalling was performed by *SMRT Link* version 9.0.0.92188 [50] followed by circular consensus sequencing (CCS) analyses to generate HiFi sequences. For Can-0 we obtained 2.9 Gbp HiFi Q20 reads with a read N50 value of 14.5 kbp, and for Col-0 1.45 Gbp HiFi Q20 reads with a read N50 value of 14 kbp.

### Genome assembly

For each accession, we first assembled the ONT and HiFi long read data separately and then merged the assemblies. Nanopore data were initially assembled by using NECAT [23] with default settings except that ‘*CNS_OUTPUT_COVERAGE*’ was changed to 40. The contigs generated by NECAT were polished by Nanopore reads using *Racon v1.4.20* [51] with options ‘*-m 8 -x -6 -g −8 -w 500’*, followed by *Medaka 1.4.4* [52] with options ‘*-m r941_prom_sup_g507*’, and finally polished with Illumina reads by *Pilon* 1.24 [53] with options *‘--changes --fix all*’. In all the polishing steps, *minimap2*[54] was used to map long-reads to the contigs, and *bwa-mem2* [55] was used to map Illumina short-read sequences. PacBio HiFi reads were assembled by *hifiasm* with option ‘-l 0’.

To combine the higher continuity from Nanopore-based contigs with the higher accuracy from PacBio HiFi-based contigs, we used *quickmerge* [56] setting the Nanopore assemblies (after three polishing steps described above) as ‘*hybrid_assembly’* and the HiFi-read assemblies as ‘*self_assembly’*, and the N50 value of the polished Nanopore-read assemblies as the minimum length cutoff (−l) of contigs to be merged. After the merging, we performed two rounds of polishing the merged assemblies with PacBio HiFi reads using racon. The alignment of HiFi reads to the merged assemblies was conducted with *pbmm2* [57]. Where the merged assembly did not retain a similar N50 value to the Nanopore-read assembly, we patched the Nanopore assembly to the merged assembly using *RagTag patch* [58]. Finally, polished merged assemblies were placed on the correct chromosomes using the published Col-0-CEN assembly [17] by using *RagTag scaffold* [58]. After scaffolding, we ran another round of final polishing of the scaffold by haplotype-aware polishing tool *Hapo-G* [59]. At each round of polishing and merging, the N50 values of assemblies or scaffolds were estimated by *QUAST* [60], the quality value (QV) of assemblies or scaffolds estimated by *Merqury* [61], and the number of Benchmarking Universal Single-Copy Orthologs (BUSCO) for estimating completeness and duplication calculated by *BUSCO* [62, 63]. Every round of polishing increased the QV of the assemblies or scaffolds.

### Omni-C sequence analysis

Genome-wide chromatin interaction data for Col-0 and Can-0 was generated using a Dovetail® (now Cantata Bio, USA) Omni-C® Kit. Plants were grown under long-day conditions for 19 d and then dark-treated for 48 h before leaf tissue was snap-frozen in liquid nitrogen. 500 mg of tissue was ground into a fine powder with liquid nitrogen using a mortar and pestle and the assay performed as per the manufacturer’s instructions for plants. Briefly, chromatin was fixed with formaldehyde and nuclease treated. An aliquot was de-crosslinked, and DNA purified with a DNA Clean & Concentrator-5 kit (Zymo Research, Irvine CA, USA). DNA yield and fragment size were determined using a Bioanalyzer high sensitivity dsDNA kit (Agilent Technologies, Milton Keynes, UK) and Qubit high sensitivity DNA quantification kit (Thermo Fisher, Altrincham, UK). The remaining lysate was then processed with reactions for end-polishing, ligation of a biotinylated oligonucleotide bridge, intra-aggregate ligation, and cross-link reversal, respectively. The DNA was purified and quantified using a Qubit high sensitivity DNA quantification kit (Thermo Fisher, Altrincham, UK) before proceeding with library preparation. Streptavidin enrichment of the biotinylated bridge was performed, and the final libraries were indexed and amplified by PCR. Illumina NovaSeq PE150 sequencing was performed by Novogene Co. Ltd (Cambridge, UK), on a Novaseq 6000 instrument. The 150 bp paired-end Omni-C reads were aligned to the merged and polished scaffolds. Read de-duplication and finding contact points was performed by following the Dovetail Omni-C kit document at https://omni-c.readthedocs.io/en/latest/index.html. *PretextMap* [64] and *PreTextView* [65] were used to generate and view the final contact map, to monitor and confirm the structure of scaffolds. Supplemental Figures S1 and S2 were generated from the contact maps using *PretextSnapshot* https://github.com/sanger-tol/PretextSnapshot [66]

### ONT methylation data generation and differential methylation analysis

ONT fast5 files were processed with *Megalodon v2.4.1*[67] and *Guppy* (GPU version *5.0.16_linux64*)[49] with the option *‘--guppy-config dna_r9.4.1_450bps_sup_prom.cfg --remora-modified-bases dna_r9.4.1_e8 sup 0.0.0 5mc CG* ‘*0*to generate the raw methylation data. The R package *NanoMethViz* [68] was used to visualize the ONT methylation data, and as well to prepare input files for *DSS* [69] for differential methylation calling between Col-0 and Can-0. The threshold P-value for calling differentially methylated loci (i.e. at individual nucleotides) was set to 0.01, and the threshold of P-value for calling differentially methylated regions (e.g. across gene bodies) was 0.05. The correlation between bisulfite and nanopore methylation results were performed by the ‘*megalodon_extras validate compare_modified_bases*’ function in the *megalodon* package.

### Comparative multi-omic analyses

Comparative assays were performed on Can-0 and Col-0 grown under long-day conditions (described above) until the emergence of the 9th rosette leaf. Leaves 3-8 were harvested and frozen in liquid nitrogen. Each biological replicate comprised five plants, pooled and homogenized by grinding with a mortar and pestle in liquid nitrogen.

### Whole-genome bisulfite sequencing (WGBS)

DNA was extracted from three biological replicates of pooled leaf tissue using a DNeasy Plant Mini Kit (Qiagen, Manchester, UK). WGBS libraries were constructed from 10 ng DNA using a Pico Methyl-Seq™ Library Prep Kit for Illumina-based Sequencing (Zymo Research, Irvine CA, USA). DNA quality control was performed using agarose gel electrophoresis and UVspectrophotometry (NanoDrop Technologies, USA). Quantification of extracted DNA was performed using a Qubit high sensitivity DNA quantification kit (Thermo Fisher, Altrincham, UK). The quality and quantity control of WGBS libraries employed an Agilent DNA 1000 Kit (Agilent technologies, Milton Keynes, UK) and Agilent 2100 Bioanalyzer. All kit workflows were performed according to manufacturers’ instructions.

Libraries were sequenced by Novagene (150 bp paired end reads, using a NovaSeq instrument). The bisulfite reads were firstly trimmed by 10 bp at each end by *trim_galore* [70], and then *bismark* [71] was run for read mapping, deduplication, and extraction of methylation information. The script *dname_bed_corr.sh*from *dna_me_pipeline* [71] was used to check the correlation of methylation between samples. Loci covered by fewer than 10 reads were removed.

### Short-read transcriptome sequencing (RNA-seq)

RNA was extracted from five biological replicates of pooled leaf tissue using a Plant RNeasy Mini Kit (Qiagen, Manchester, UK) as per the manufacturer’s recommendations. DNase treatment was performed using a TURBO DNA-free kit (Invitrogen, now ThermoFisher Scientific). RNA quantification and analysis of integrity employed an Agilent RNA 6000 Nano Kit (Agilent technologies, Milton Keynes, UK) and the Agilent 2100 Bioanalyzer. Illumina NovaSeq PE150 sequencing was performed by Novogene Co. Ltd, UK. The sequencing data were trimmed and filtered by *BBduk* [48] with options ‘*qtrim=rl trimq=10 maq=10*’ to achieve clean and good quality sequences. mRNA expression levels were summed across all isoforms for a given gene using *kallisto* [72] to produce normalized transcripts per million (TPM) values within each replicate; the expression of a gene across replicates was estimated by the log_10_ of their geometric mean, after adding a pseudocount of 1.0 to avoid negative infinite values.

### tRNA quantification (mim-tRNAseq)

RNA was isolated from two biological replicates of pooled leaf tissue using phenol/chloroform extraction. Tissue was cryo-pulverised in liquid nitrogen using a Geno/Grinder® (Spex SamplePrep 2010 USA) and 1 ML of of TRIzol™ Reagent was added to 300-400 μL tissue powder, followed by incubation at RT for 5 min. RNA was extracted by the addition of 0.2 vol chloroform, incubation at RT for 2 min and centrifugation at 12,000 g for 15 min at 4°C. The upper aqueous phase was extracted with an equal volume of chloroform and the RNA precipitated by addition of ice-cold 100 % ethanol. Following centrifugation at 12,000 g for 20 min at 4°C, the pellet was washed in 80 % (v/v) ethanol, air-dried, and resuspended in RNAse-free water. RNAs were sequenced using modification-induced misincorporation tRNA sequencing (mim-tRNAseq)[32, 33]

Data analysis was performed using the bioinformatics pipeline in [32]. In brief, the sequences were trimmed with *cutadapt* version 4.1[73]; a first step trims the GATATCGTCAAGATCGGAAGAGC adapter in 31, a second step trims the two bases due to the circularization *(−u 2*), and a last run cuts the remaining adapter in 51: CTTGACGATATC). Only reads longer than 25 bp with quality >25 were retained for mim-tRNAseq analysis. The package *mim-tRNAseq* version 1.1.7 [32] was used with the parameters ‘*–species Atha–cluster-id 0.95–threads 15–min-cov 0.0005–max-multi 4–remap–remap-mismatches 0.075*’. We determined the Araport11 ids of the resulting tRNA genes based matching their TAIR10 coordinates.

### Ribosome profiling

Ribonucleic complexes from each accession were solubilised from six biological replicates of pooled leaf tissue and clarified as per [74]. Polysomes were separated on a sucrose gradient with absorbance measured at 254 nm using a UV-1 monitor (Pharmacia, Uppsala, Sweden). Ribosome profiling (3’Riboseq) was performed as described in [75] with pooling of monosome and polysomes. Ribosome-associated RNA was precipitated using sodium acetate and ethanol and purified using a Zymo quick RNA column (Zymo Research, Irvine CA) and the integrity assessed using an Agilent Bioanalyzer 2100. mRNA libraries were prepared by Novogene Co. Ltd., UK and sequenced using NovaSeq to produce paired end 150 bp reads. Data was analysed as for RNA-seq, with additional filtering to remove ribosomal RNAs and tRNAs using sequences from [76].

### Quantitative proteomics

#### Protein extraction, reduction, alkylation, and digestion

Total protein was extracted from four biological replicates of pooled leaf tissue. Tissue samples (100 mg) were cryo-pulverised in liquid nitrogen using a mortar and pestle. Protein was precipitated by the addition of 5 mL pre-chilled 10% (w/v) trichloroacetic acid (TCA) in acetone, followed by incubation at −20 °C overnight. The precipitated protein pellet was washed three times with chilled acetone. TCA-precipitated pellets were solubilised and reduced in 100 μL of urea containing buffer [8 M urea, 50 mM triethylammonium bicarbonate (TEAB), 10 mM dithiothreitol, 1x protease inhibitor cocktail (Roche, Mannheim, Germany)] at 25 °C for 1 hour. Protein was alkylated by the addition of 2-chloroacetamide to achieve a final concentration of 55 mM, followed by incubation at 25 °C for 30 minutes. In-solution protein digestion was performed in three sequential steps: for the first digest, 2 μg of Lys-C (Promega, Madison, WI) was added to give a 1:100, w/w enzyme-protein ratio, followed by incubation at 37 °C for 4 hours. For the second digest, 700 μL of 50 mM TEAB was added to reduce the urea concentration below 1 M, and 2 μg of trypsin (Promega) was added. The mixture was incubated at 37 °C overnight, followed by the addition of a further 2 μg of trypsin for the third digest and a 4-hour incubation at 37 °C. The digests were acidified with 1% trifluoroacetic acid (TFA) and desalted using 50 mg SepPak tC18 cartridges (Waters Corporation, Borehamwood, UK). The cartridge was washed with 0.1% TFA solution and eluted in two steps: (i) 300 μL 25% acetonitrile (ACN) in 0.1% formic acid (FA); (ii) twice with 300 μL 50% ACN in 0.1% FA. The eluates were lyophilised and stored at −20 °C. Peptide amount was determined using a Pierce Quantitative Colorimetric Peptide Assay kit (Thermo Fisher Scientific, Hemel Hempstead, UK).

#### Mass spectrometry

The proteomics data was acquired using a timsTOF HT mass spectrometer (Bruker Daltonics, Bremen, Germany) coupled with a nanoElute 2 UHPLC system (Bruker Daltonics). Peptides (750 ng) were loaded onto a PepMap Neo trap column (300 µm x 5 mm, 5 µm particle size, Thermo Scientific) and separated on a µPAC Neo analytical column (500 mm x 180 μm, 16 μm pillar length, Thermo Scientific) using a 60-min non-linear gradient consisting of 5%–17% solvent B over 42 min at a flow rate of 300 nL/min, followed by an increase to 26% for 14 min and 37% for 4 min. The mobile phases comprised 0.1% FA in water as solvent A and 0.1% FA in ACN as solvent B. The eluates were ionised using a Captive Spray source via a ZDV Sprayer emitter (20 µm, Bruker Daltonics). The mass spectrometer was set to dia-PASEF scan mode spanning 100–1700 m/z in positive ion mode. The ion mobility (IM) range was set to 0.85–1.23 1/K0 [V s/cm2], and both the ramp time and the accumulation time was set to 100 ms, corresponding to a ramp rate of 9.42 Hz. The variable collision energy was applied depending on the IM, ranging from 20 eV at 0.60 1/K0 to 59 eV at 1.6 1/K0. The dia-PASEF windows were optimised for the *Arabidopsis* proteome profile using *py_diAID* version 0.0.19 [77]. Ten dia-PASEF scans were divided into 3 IM windows with a mass range of 300–1,200 Da, corresponding to an estimated cycle time of 1.17 s.

#### Proteome Data analysis

The mass spectra from Col-0 and Can-0 were searched separately against their own sequence databases using *DIA-NN v1.8.2 beta 27* [78] The Col-0 and Can-0 protein sequence databases derived from *de novo* assemblies were used to generate *in silico* spectral libraries, which contain predicted retention times and predicted ion mobility (1/K0) values. A maximum of one missed cleavage was permitted with a minimum peptide length of 7 amino acids. Dynamic modifications considered oxidised methionine, acetylation at the protein N-terminus, and methionine loss at the protein N-terminus. Carbamidomethylation of cysteine was designated as a fixed modification. The *’match between runs*’ option was utilised to minimise missing identifications. The R package *DIAgui* version *1.4.2* [79] was used to generate iBAQ values [80] from the protein intensity, filtered at both precursor and gene levels at 1% FDR, using only proteotypic peptides. The protein matrices from Col-0 and Can-0 were combined using the Hierarchical Orthologous Group (HOG) classification, which identifies orthologous genes between accessions (see below). The iBAQ value for gene *i* was normalized using the equation:

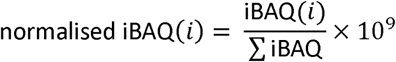

### Long-read transcriptome sequencing (Iso-seq)

Iso-seq data were generated from Can-0 and Col-0 rosette leaves grown under long-day conditions (described above) to support genome annotation only. Total RNA was extracted from one biological replicate of pooled leaf tissue, using a Plant RNeasy Mini Kit (Qiagen, Manchester, UK) as per the manufacturer’s recommendations. Quantification of RNA and analysis of integrity was performed using an Agilent RNA 6000 Nano Kit (Agilent Technologies, Stockport UK). PacBio Iso-Seq™ SMRTbell® libraries were constructed by Novogene Co. Ltd, UK with the Express Template Prep Kit 2.0 for sequencing on Sequel® II System. Sequencing data were analysed according to the *Isoseq3* instructions[81], including consensus sequence generation by *ccs* [82], primer removal and demultiplexing by *lima*, trimming the polyA tail and concatemer by *isoseq*. The clustering step suggested by the Isoseq3 workflow was omitted.

### Genome annotation

The finalised genome assemblies of Col-0 and Can-0 were first scanned by *EDTA* [83] with options ‘*-- sensitive 1 --evaluate 1 --anno 1*’ to mask and annotate repeats, using the *RepBase*[84] transposable element (TE) database (version 27.01) of *A. thaliana* as the curated TE library and the known coding sequences of *A. thaliana* (Araport11 [19]). The sequences of annotated repeats were aligned against the Araport11 coding sequences using *blastn* [85], to ensure that no gene sequences overlapped with the repeats identified by *EDTA*.

After cleaning and QC, the Illumina RNAseq reads for Col-0 and Can-0 were each aligned to their respective assembled genomes by *STAR* [86] with option *‘--outSAMstrandField intronMotif*’ to generate RNAseq alignment *.bam* files. The *.bam* files together with the soft-masked genomes generated by *EDTA* were then input to *Braker1* [87] for gene prediction and annotation. We used the option *‘--UTR=on --augustus_args="--species=arabidops*"*is*’ to apply the *Augustus* [88] pre-trained gene model of *Arabidopsis* for *ab initio* gene prediction and UTR annotation. We used *Braker2* to annotate genes using the Uniprot *A.thaliana* proteome database [89], and the long-read protocol from *Braker* to generate a version of annotation that incorporated our PacBio Isoseq data. We mapped the filtered Iso-Seq reads to the genome by *minimap2* [90] with option *‘-ax splice -uf -- secondary=no -C5*’ following the long-read protocol from *Braker* [91]. The three gene annotations respectively from *Braker1*, *Braker2*, and long-read protocol were then merged and filtered by *Tsebra* [92] with the option ‘*long_reads_filtered.cfg*.*’*

We used the *PASA* pipeline [93] to refine these *Tsebra* annotations. First *Trinity* [94] produced genome-guided and *de novo* assembled transcriptomes from both Isoseq and Illumina RNAseq. Secondly, we ran the *PASA* pipeline using default options with two rounds of annotation updates on the Tsebra annotation. To make sure that we have the most complete gene set for the accessions, we also augmented our updated annotations with the lifted over annotation from Tair10 which was produced by *Liftoff* [95] with default settings. This final step was performed by the function *agat_sp_complement_annotations.pl* from *AGAT* v.0.9.2 [96] using the *PASA* updated annotation as the reference and adding any additional genes from lifted over annotations. Functional annotation for the finalised annotations was conducted by *InterProScan* [97] on https://usegalaxy.eu [98]. We used *AGAT* to integrate functional and homology annotations into GFF format. Finally, we used *BUSCO* to estimate the completeness of the annotated transcriptomes, and *AGAT* to correct small problems in the annotation such as duplication of genes with different identifiers and changing gene IDs.

We identified orthologous genes using *OMA standalone* [99]. We downloaded the OMA database orthologs from the five Brassica species (*Arabis alpina*, *Arabidopsis lyrate*, *Arabidopsis thaliana (Tair10)*, *Brassica napus*, *Brassica oleracea*, *Brassica rapa subsp. Pekinensis*) and combined them with the TAIR10, Col-0 and Can-0 protein sequences into Hierarchical Orthologous Groups (HOG) using *pyHam* [100]. For HOGs containing more than one gene from the same accession, gene DNA sequences were used to identify the most similar homologs inter-accessions by reciprocal *blastn* [85, 101] searches. If were more than two genes from each same accession in a HOG, we used DNA sequence of each gene to search for the genes pairs between accessions that are the most similar, to find the 1-to-1 homologous genes.

### Comparative genomics

Our assemblies of Col-0 and Can-0 were aligned using *minimap2* [90]. Synteny statistics were calculated *by dna_diff* [20]. High-confidence variants including indels below 10bp were called by *clair3* [102] and structural variants were called using *pbsv* [103]with HiFi reads.

### Differential expression analysis

Clean RNAseq data from Col-0 and Can-0 were fed into *Trinity v2.14.0* [94] for RNAseq quantification and differential expression analysis, using default parameters with Col-0 as the reference genome. We used *kallisto* [72] for pseudo-alignment of RNAseq reads and quantification of gene and isoform expression. The output from *kallisto* was used as input for *EdgeR* [104] for normalization and differential expression analysis. The gene expression within each replicate sample was normalized by calculating transcripts per million (TPM), and inter-sample normalization was performed by calculating Trimmed Mean of M-values (TMM) in *EdgeR*. We used a FDR threshold of 0.05 to identify genes or transcripts that are differentially expressed between Col-0 and Can-0.

The normalised iBAQ proteome values were log-transformed and missing values imputed with random values from a normal distribution (width 0.3, down shift 1.8) using *Perseus v1.6.15.0* [105]. DE proteins were determined using Student t-test with threshold of 0.05 for the Benjamini-Hochberg adjusted p-value (FDR) and a 2-fold change.

### Testing relationships between annotations and expression

Relationships between a dichotomous annotation difference between Col-0 and Can-0 (such as differential gbM or the presence/absence of a TE upstream of a gene) and continuous mRNA or protein expression were tested by first subdividing the genes into two subsets corresponding to those genes with and without the specified attribute (eg whether or not the gene is differentially methylated) and then testing if the mean of the absolute values of the log_2_ fold-change of mRNA or protein expression in the two subsets differed, using a non-parametric Mann-Whitney test implemented in R. Statistical significance was reported as logP, the negative log_10_(p-value) of the test. This methodology does not assume any particular direction of effect between annotation and expression.

## Supplemental Figures

***Supplemental Figure S1.*** *Omni-C contact map as generated by PretextSnapshot for Col-0*

***Supplemental Figure S2.*** *Omni-C contact map as generated by PretextSnapshot for Can-0*

***Supplemental Figure S3.*** *Distribution of mRNA and protein expression restricted to genes expressed in all replicates, scaled so that the median level of expression of genes with both protein and mRNA expression is equal to 1. A, B: scatter plots of mRNA (x-axis) vs protein (y-axis) expression for orthologous genes in Col-0 (A) and Can-0 (B). Dotted red lines show medians. C,D: Histograms of mRNA expression for genes with (pale red) or without (grey) detectable protein expression in Col-0 (C) and Can-0 (D). The black curves indicate lognormal densities fitted to the mRNA+protein histograms using robust estimates of mean and standard deviation. Expression scales are logarithmic throughout.*

***Supplemental Figure S4.*** *Heatmaps of CpG methylation correlation between methylation estimated from bisulphite-converted Illumina vs Oxford Nanopore sequence. (A) 2.6M CpG sites in Col-0 (B) 2.8M CpG sites in Can-0.*

***Supplemental Figure S5.*** *Estimated codon effects on mRNA and protein expression in Col-0. Shown are barplots representing the multiple regression coefficients for 61 non-termination codons (y-axis) for modelling log-transformed mRNA (red) or protein expression (blue). Error bars show standard errors.*

***Supplemental Figure S6.*** *Scatter plots of the estimated codon expression effects for mRNA (A,B) or protein (C,D) vs tRNA abundance in Col-0 (A,C) and Can-0 (B,D). Each dot represents one codon, labelled by its codon and encoded amino acid. The black numbers are the Pearson correlation coefficients.*

***Supplemental Figure S7.*** *Differentially expressed transcripts. Of 17,771 transcripts quantified, 7.585 were differentially expressed (FDR <0.05). A. Volcano plot showing the relationship between statistical significance (adjusted p-value) on the y-axis and the biological significance (log2 fold change) on the x-axis. B. Gene ontology terms enrichment for different groups of differentially expressed genes (DEGs).*

***Supplemental Figure S8.*** *Differentially expressed proteins. Of 8915 proteins quantified, 1196 were differentially expressed (>2-fold-change, adj. p <0.05). A. Volcano plot showing the relationship between statistical significance (adjusted p-value) on the y-axis and the biological significance (log2 fold change) on the x-axis. B. Gene ontology terms enrichment for different groups of differentially expressed proteins (DEPs).*

## Supplemental Tables

***Supplemental Table S1*** *Summary statistics for our assemblies of Col-0 and Can-0, and comparisons with three published Col-0 assemblies. Base QV estimates the error rate in the assembly, expressed as the negative log10 of the probability a given base pair is erroneous* [61]*, BUSCO estimates the completeness of the gene content of the assembly in terms of 4596 single-copy orthologs found in brassicas* [62, 63]*. Assembly N50 is the length of contigs such that 50% of the assembly is in contigs of at least N50. Scaffold N50 is the corresponding length in assembled scaffolds. #Gaps is the number of gaps in the assembly. GC content is the percentage of G+C nucleotides in the assembly. Genome size is the total length of the assembly.*

***Supplemental Table S2*** *Counts of differences as computed by dna_diff* [20] *between our Can-0 (blue background) and Col-0 (yellow background() assemblies and with four other Col-0 and one other Can-0 assemblies, namely Can-Lian, Col-Lian:* [16]*, Col-CEN* [17] *, Col-XJTU* [18] *, Col-TAIR, the TAIR10 reference, Col-CC: the community consensus assembly (Genbank id GCA_028009825.2). Pink background shows comparisons between selected other Col-0 assemblies. The data are shown graphically in Figure 2.*

***Supplemental Table S3*** *Orthologous genes between Col-0, Can-0 and Araport11. Each row represents one Homologous Group (HOG). Numbers of alternatively spliced isoforms for each gene are indicated by the “Transcripts” columns.*

***Supplemental Table S4*** *Normalised expression values for gbM, mRNA, protein and ribo-RNA. Values are supplied for each replicate (except for gbM) and combined across replicates, and log-transformed in both accessions Col-0 and Can-0.*

***Supplemental Table S5*** *Pearson correlations between replicates for log-transformed mRNA, protein and ribo-RNA expression values. Light blue background indicates correlations between replicates within an accession, white background indicates correlations between replicates from different accessions*

***Supplemental Table S6*** *tRNA expression values for Col-0 and Can-0.*

***Supplemental Table S7*** *Estimated codon effects in Col-0 and Can-0 together with the tRNA codon effects*

***Supplemental Table S8*** *ANOVA tables used to generate Figure 9. Summary of omics multiple regression models, relating to the barplots the Figure. Each block of consecutive rows with the same background shade share the same Model specification^a^. Each row describes the variance component listed as the source of variation ^b^. The degrees of freedom for the component in the analyses of variance is in column df ^c^. The negative base-10 logarithm of the p-value for the variance component is in column logP ^d^. The percentage of variance explained by the component after fitting the preceding components in the model is in column R^2^%^e^ (these are the values displayed in the stacked barplots in Figure 9). The cumulative percentage of variance explained by the components is in column Cum R^2^ %^f^. Each model is in the format Y ∼ X, where Y is the dependent variable and X is one or more independent explanatory variables. The dependent variables for the Col-0 analyses are:- Col-protein, log-transformed protein expression; Col-mRNA: log-transformed mRNA expression for genes also with protein expression; Col-mRNA*: log-transformed mRNA expression for genes without protein expression; Col-gbM: percent gene-body methylation for genes also with protein expression; Col-gbM*: percent gene-body methylation for genes without protein expression. Similar names apply for the Can-0 analyses. The independent explanatory variables are:- CDS: DNA sequence composition; gbM: percent gene body methylation; mRNA: log-transformed mRNA expression. CDS effects are subdivided into protein.length, amino.acid.usage and codon.usage;*

***Supplemental Table S9*** *Differential Expression analysis between Col-0 and Can-0. Workbook DE mRNA shows the EdgeR analysis output for mRNA. Workbook DE protein shows the analysis output for protein.*

