## Supplementary material for "Revisiting the Central Dogma: the distinct roles of genome, methylation, transcription, and translation on protein expression in *Arabidopsis thaliana*": Compressed archive of all supplemental files: Supplemental Figure S7 differential expressed mRNA and protein.pptx (from another version of this preprint: version 1, 2025-01-10)

### Slide 1
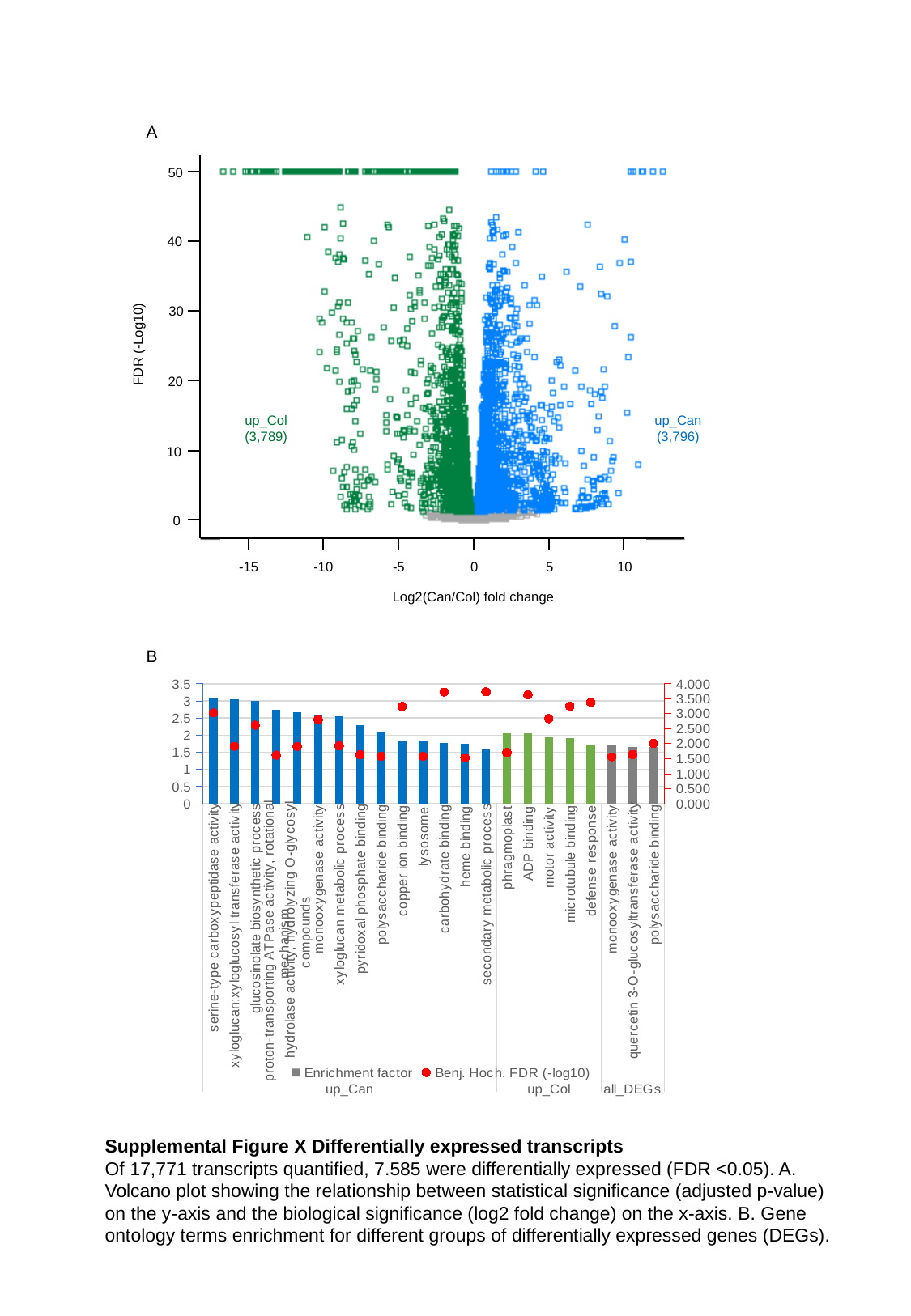

A
50
40
30
FDR (-Log10)
20
up_Col
(3,789)
up_Can
(3,796)
10
0
-15
-10
-5
0
5
10
Log2(Can/Col) fold change
B
#### Chart
| Category | Enrichment factor | Benj. Hoch. FDR (-log10) |
|---|---|---|
| serine-type carboxypeptidase activity | 3.061 | 3.0271401980412334 |
| xyloglucan:xyloglucosyl transferase activity | 3.043 | 1.914531030113328 |
| glucosinolate biosynthetic process | 2.9962 | 2.619228156559993 |
| proton-transporting ATPase activity, rotational mechanism | 2.7309 | 1.6199516492630748 |
| hydrolase activity, hydrolyzing O-glycosyl compounds | 2.6751 | 1.9096713931765776 |
| monooxygenase activity | 2.5748 | 2.8033541132129125 |
| xyloglucan metabolic process | 2.5535 | 1.9308871486128791 |
| pyridoxal phosphate binding | 2.2837 | 1.6336891332332644 |
| polysaccharide binding | 2.0807 | 1.5828605902726745 |
| copper ion binding | 1.8453 | 3.244712219460692 |
| lysosome | 1.8392 | 1.5823788621361057 |
| carbohydrate binding | 1.7834 | 3.7232166152997315 |
| heme binding | 1.7441 | 1.5298248033448205 |
| secondary metabolic process | 1.587 | 3.7284604144061966 |
| phragmoplast | 2.0608 | 1.7044548901882588 |
| ADP binding | 2.0571 | 3.628913657505206 |
| motor activity | 1.9408 | 2.83612334435308 |
| microtubule binding | 1.9136 | 3.249576466712681 |
| defense response | 1.7328 | 3.385968017798962 |
| monooxygenase activity | 1.6986 | 1.5634239360009399 |
| quercetin 3-O-glucosyltransferase activity | 1.6661 | 1.6384043606817655 |
| polysaccharide binding | 1.6487 | 2.0121461174960045 |Supplemental Figure X Differentially expressed transcripts
Of 17,771 transcripts quantified, 7.585 were differentially expressed (FDR <0.05). A. Volcano plot showing the relationship between statistical significance (adjusted p-value) on the y-axis and the biological significance (log2 fold change) on the x-axis. B. Gene ontology terms enrichment for different groups of differentially expressed genes (DEGs).

### Slide 2
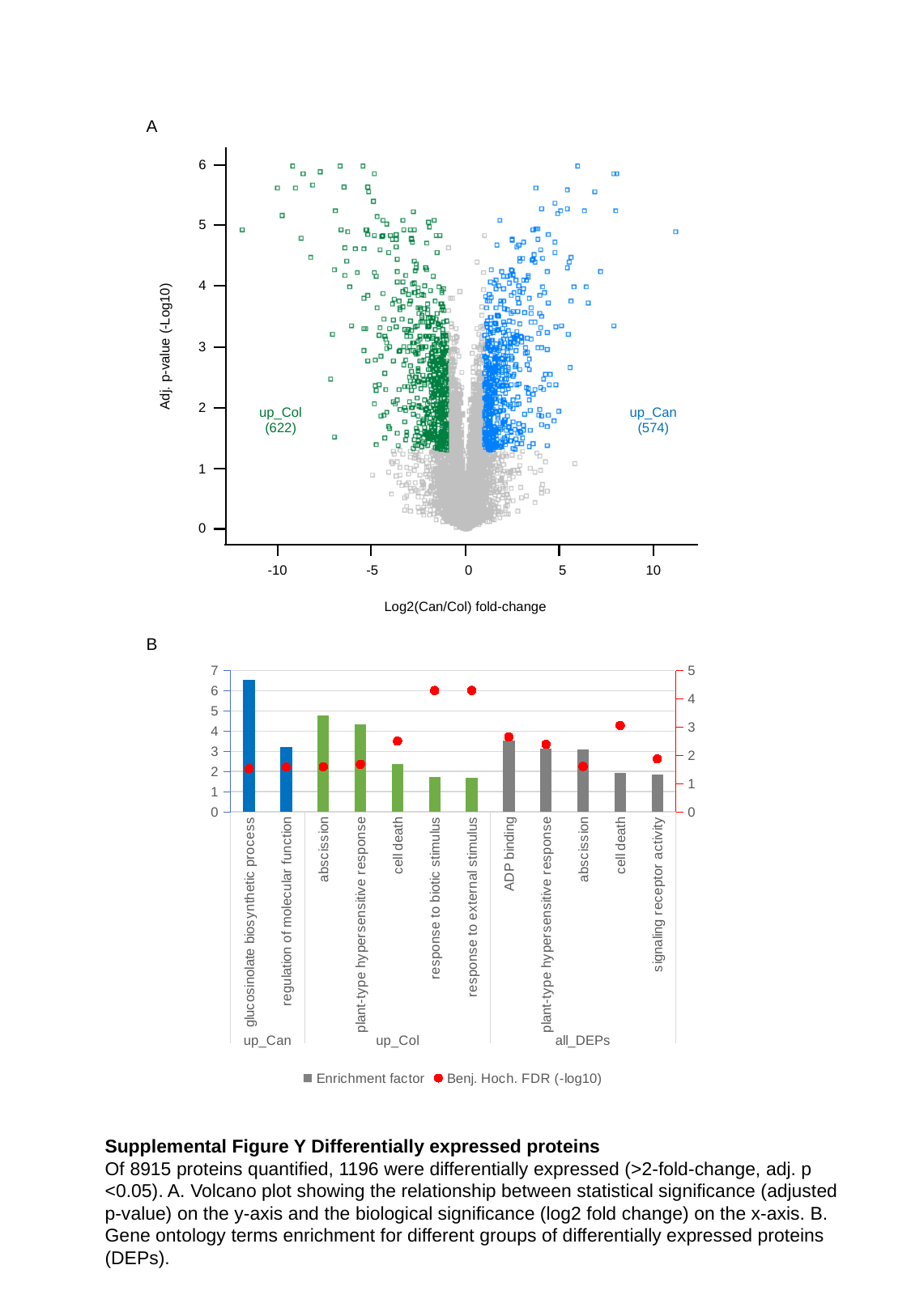

A
6
5
4
3
Adj. p-value (-Log10)
2
up_Col
(622)
up_Can
(574)
1
0
-10
-5
0
5
10
Log2(Can/Col) fold-change
B
#### Chart
| Category | Enrichment factor | Benj. Hoch. FDR (-log10) |
|---|---|---|
| glucosinolate biosynthetic process | 6.5395 | 1.5259878148301498 |
| regulation of molecular function | 3.2134 | 1.584676017300308 |
| abscission | 4.7776 | 1.5956110644694559 |
| plant-type hypersensitive response | 4.3332 | 1.684911726911683 |
| cell death | 2.3888 | 2.50020298332661 |
| response to biotic stimulus | 1.7282 | 4.294735137682596 |
| response to external stimulus | 1.6862 | 4.294735137682596 |
| ADP binding | 3.52 | 2.654410646259351 |
| plant-type hypersensitive response | 3.1203 | 2.388372268531713 |
| abscission | 3.1058 | 1.6061792046591326 |
| cell death | 1.9325 | 3.057590100340485 |
| signaling receptor activity | 1.8635 | 1.8741610360049958 |Supplemental Figure Y Differentially expressed proteins
Of 8915 proteins quantified, 1196 were differentially expressed (>2-fold-change, adj. p <0.05). A. Volcano plot showing the relationship between statistical significance (adjusted p-value) on the y-axis and the biological significance (log2 fold change) on the x-axis. B. Gene ontology terms enrichment for different groups of differentially expressed proteins (DEPs).
