## Supplementary material for "Revisiting the Central Dogma: the distinct roles of genome, methylation, transcription, and translation on protein expression in *Arabidopsis thaliana*": Compressed archive of all supplemental files: Supplemental File S7.docx (from another version of this preprint: version 1, 2025-01-10)

Supplemental File XX

***Overlap between codon effects on expression with mRNA half-life data.***

In the study (Agarwal V, Kelley DR: **The genetic and biochemical determinants of mRNA degradation rates in mammals.** *Genome Biol* 2022, **23:**245), Figure 3C lists codons and other genomic features that affect mRNA half lives. The relevant figure is reproduced here:

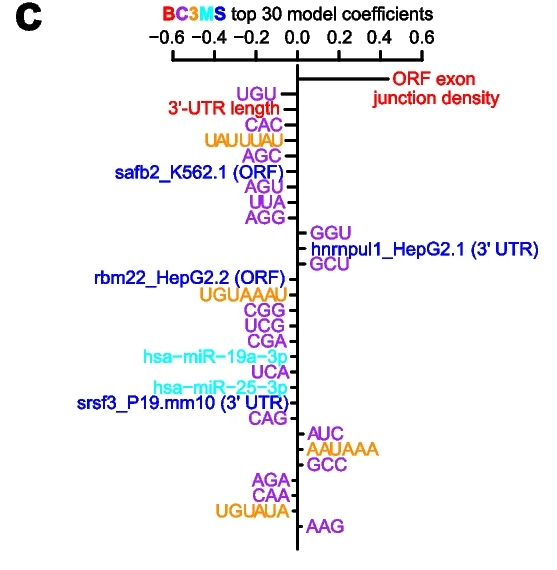

The figure shows the top 30 ranked feature coefficients. The sign of a feature’s effect on half life is indicates on the x-axis (positive: right, negative: left). The table below lists the signs of the codons taken from the above figure and the signs of the codon effects on mRNA and protein abundance in our study (Supplemental Table .

| Codon | Argarwal sign | This study sign | Agreement |
| --- | --- | --- | --- |
| UGU | - | - | TRUE |
| CAC | - | - | TRUE |
| AGC | - | - | TRUE |
| AGU | - | - | TRUE |
| UUA | - | - | TRUE |
| AGG | - | - | TRUE |
| GGU | + | + | TRUE |
| GCU | + | + | TRUE |
| CGG | - | - | TRUE |
| UCG | - | - | TRUE |
| CGA | - | - | TRUE |
| UCA | - | - | TRUE |
| CAG | - | + | FALSE |
| AUC | + | 0/+ | UNCLEAR |
| GCC | + | + | TRUE |
| ACA | - | - | TRUE |
| CAA | - | - | TRUE |
| AAG | + | + | TRUE |

For 16/18 codons there is clear agreement, for codon AUC the result is unclear and for codon CAG there is disagreement. The fraction of positive coefficients in Agarwal is $p=5/18=0.2778$, and the fraction in our study is also $p=5/18$ if we interpret AUC as 0. If the allocation of signs to codons is independent in the two studies, then the probability that a codon has the same sign is $q= p^{2}+ {(1-p)}^{2} = 0.5987$. The probability of observing a given number of agreements is from the binomial distribution $B(18, q)$. The probability of 16 or more agreements is therefore given by the R function call

pbinom(15,18,prob= 0.5987654,lower=FALSE) = 0.008.
