## Supplementary material for "Revisiting the Central Dogma: the distinct roles of genome, methylation, transcription, and translation on protein expression in *Arabidopsis thaliana*": Compressed archive of all supplemental files: Supplemental Table S1.docx (from another version of this preprint: version 1, 2025-01-10)

|  |  | **Col-0** | **Can-0** | **Col_CEN** | **Col_XJTU** | **Col_Tair10** |
| --- | --- | --- | --- | --- | --- | --- |
| **Size (bp)** | **Chr1** | 32641497 | 34797179 | 32540122 | 32659241 | 30427671 |
|  | **Chr2** | 22816092 | 23408526 | 22217084 | 22560461 | 19698289 |
|  | **Chr3** | 25863195 | 24971814 | 25743512 | 26161332 | 23459830 |
|  | **Chr4** | 21559647 | 19452999 | 21578073 | 22250686 | 18585056 |
|  | **Chr5** | 29666167 | 30003755 | 29480885 | 30093473 | 26975502 |
| **Base QV** | **Chr1** | 58.84 | 55.65 | 61.44 | 67.78 | 48.46 |
|  | **Chr2** | 59.95 | 58.03 | 45.57 | 61.89 | 52.3 |
|  | **Chr3** | 51.17 | 60.07 | 57.52 | 66.16 | 51.27 |
|  | **Chr4** | 64.76 | 57.12 | 49.93 | 66.73 | 44.7 |
|  | **Chr5** | 52.61 | 61.06 | 51.36 | 63.95 | 48.76 |
| **BUSCO** | **Single** | 4501 (97.9%) | 4500 (97.9%) | 4503 (98.0%) | 4503 (98.0%) | 4503 (98.0%) |
|  | **Duplicate** | 59 (1.3%) | 62 (1.3%) | 59 (1.3%) | 59 (1.3%) | 58 (1.3%) |
|  | **Fragmented** | 3 (0.1%) | 3 (0.1%) | 2 (0.0%) | 2 (0.0%) | 3 (0.1%) |
|  | **Missing** | 33 (0.7%) | 31 (0.7%) | 32 (0.7%) | 32 (0.7%) | 32 (0.6%) |
| **Assembly N50** | | 18,399,411 | 21,470,873 | - | 22,250,686 | 11,194,537 |
| **Scaffold N50** | | 25,863,195 | 24,972,360 | - | 26,161,332 | 23,459,830 |
| **#Gaps** | | 17 | 5 | - | 2 | 93 |
| **GC content** | | 36.28% | 36.31% | - | 36.34% | 36.03% |
| **Genome size** | | 133,233,343 | 133,091,828 | 131,559,676 | 133,725,193 | 119,668,634 |
