## Supplementary figures and images for "Revisiting the Central Dogma: the distinct roles of genome, methylation, transcription, and translation on protein expression in *Arabidopsis thaliana*" (from another version of this preprint: version 1, 2025-01-10)

### Supplemental Figure S1.png

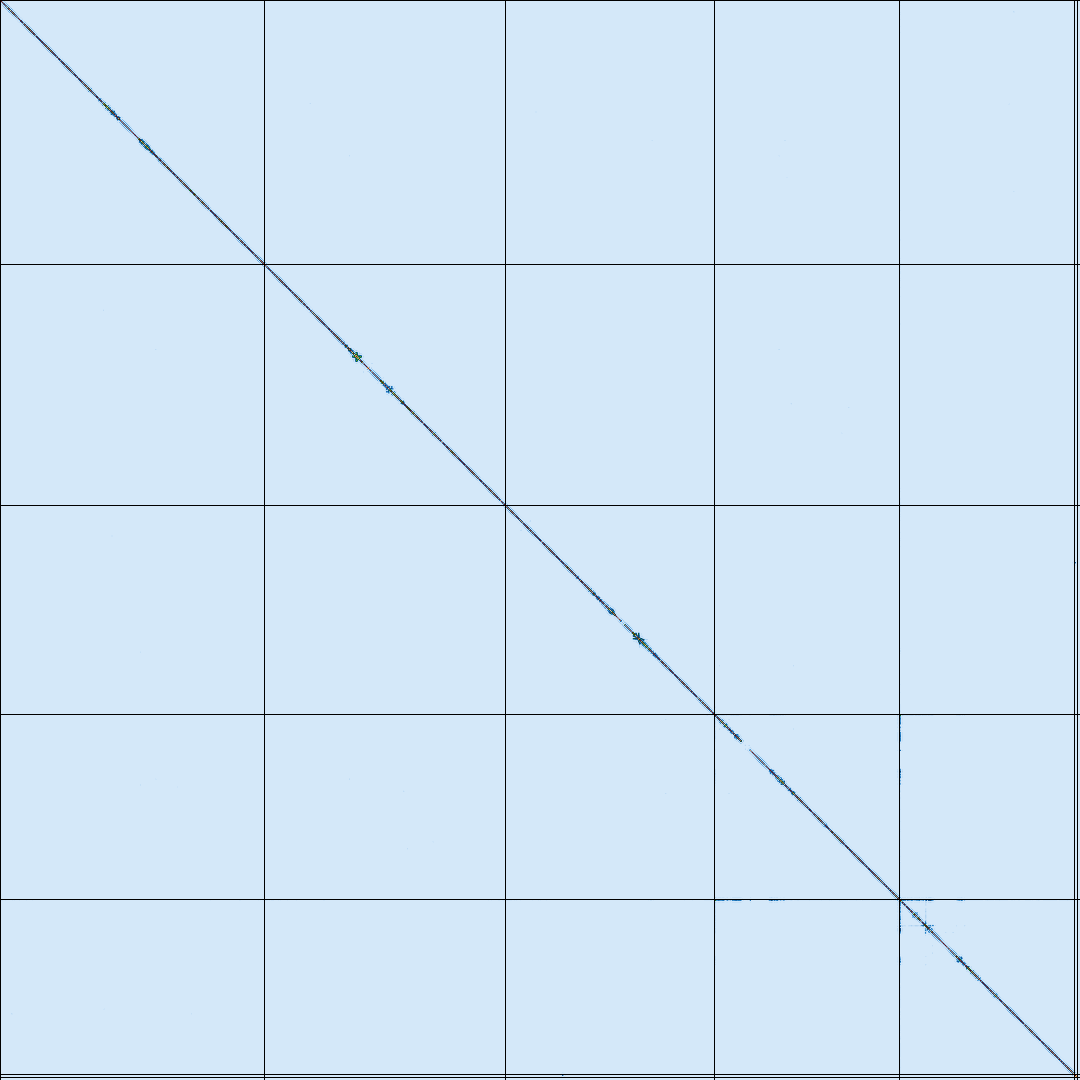

### Supplemental Figure S2.png

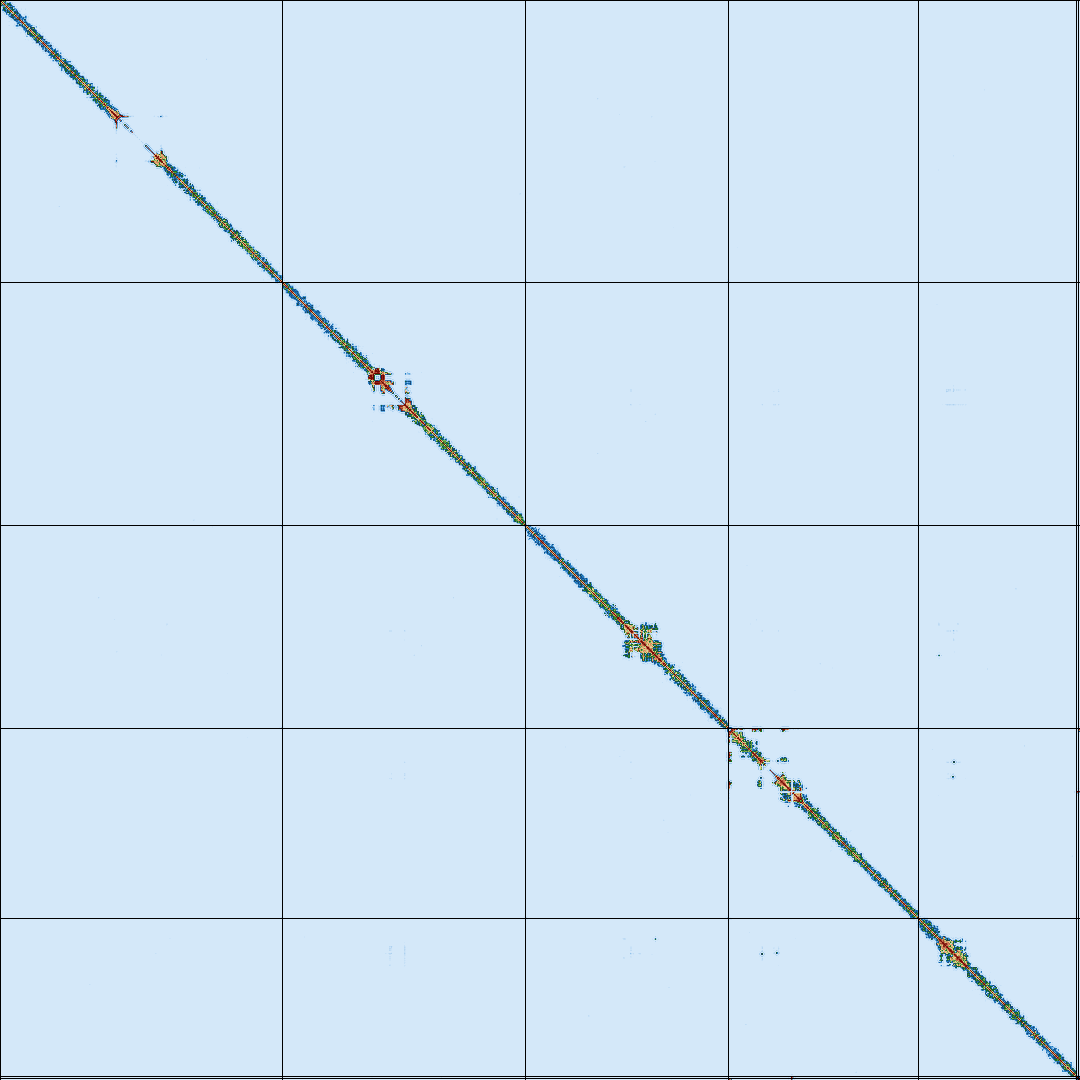

### Supplemental Figure S3.png

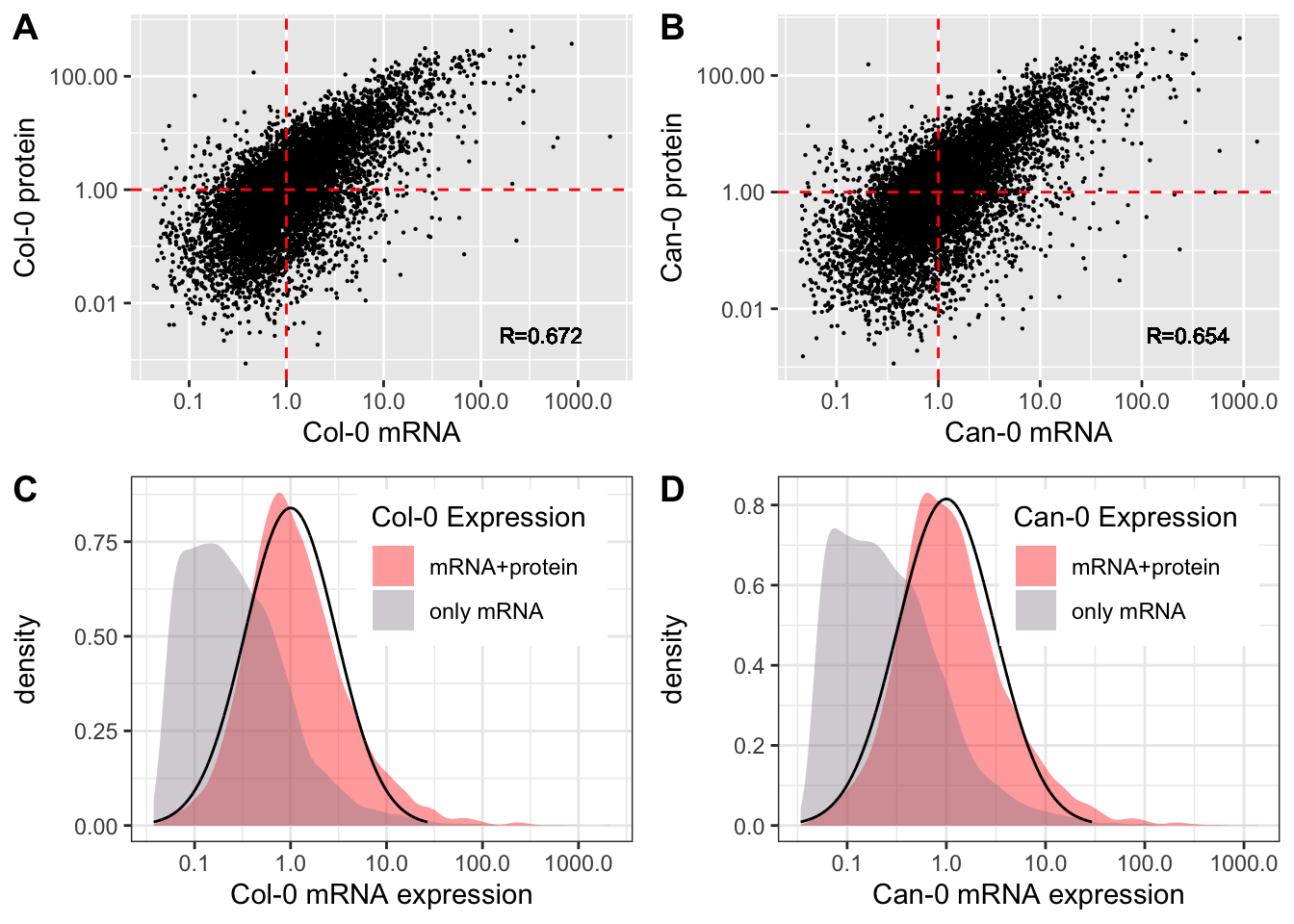

### Supplemental Figure S5.png

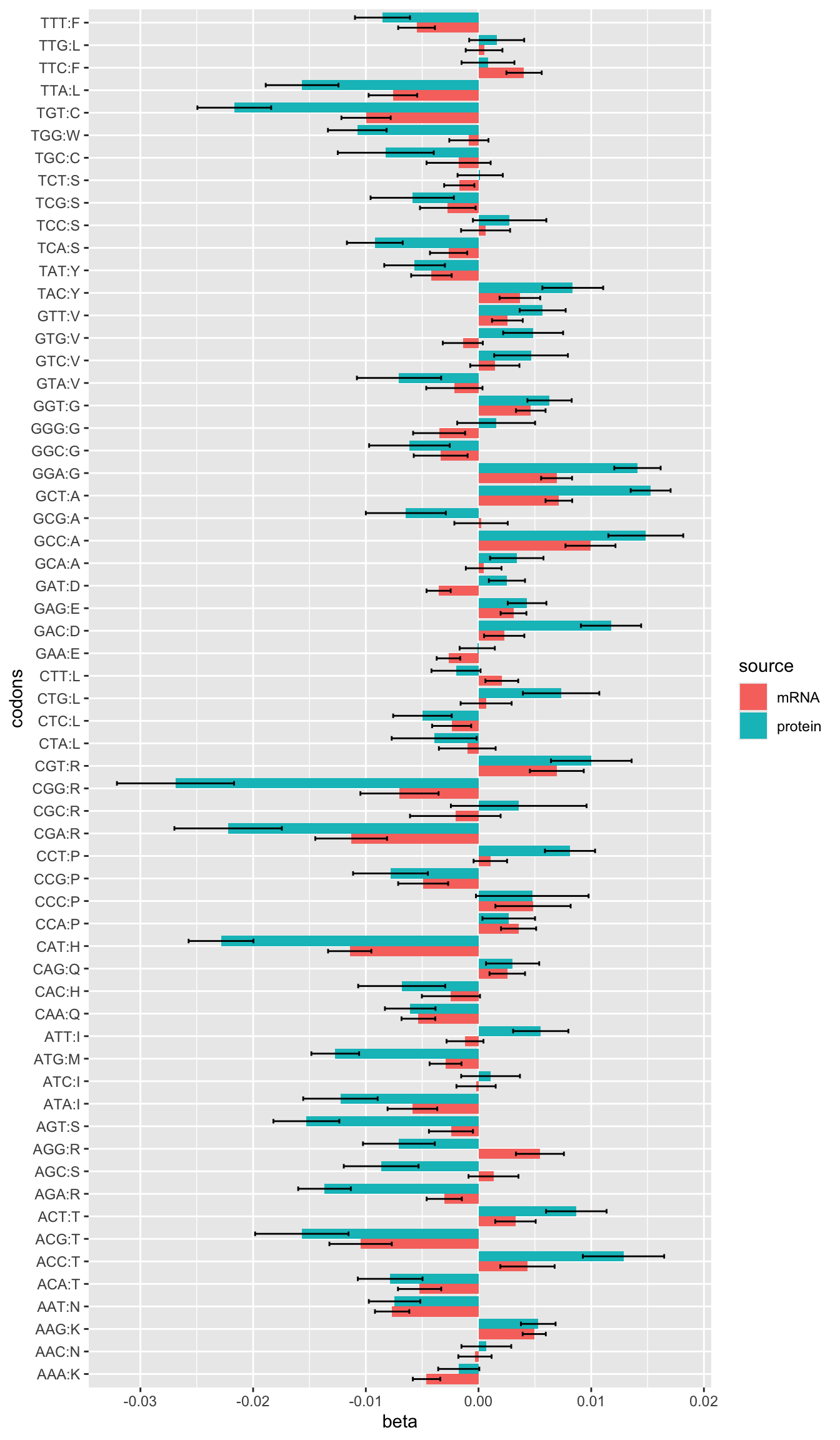

### Supplemental Figure S6.png

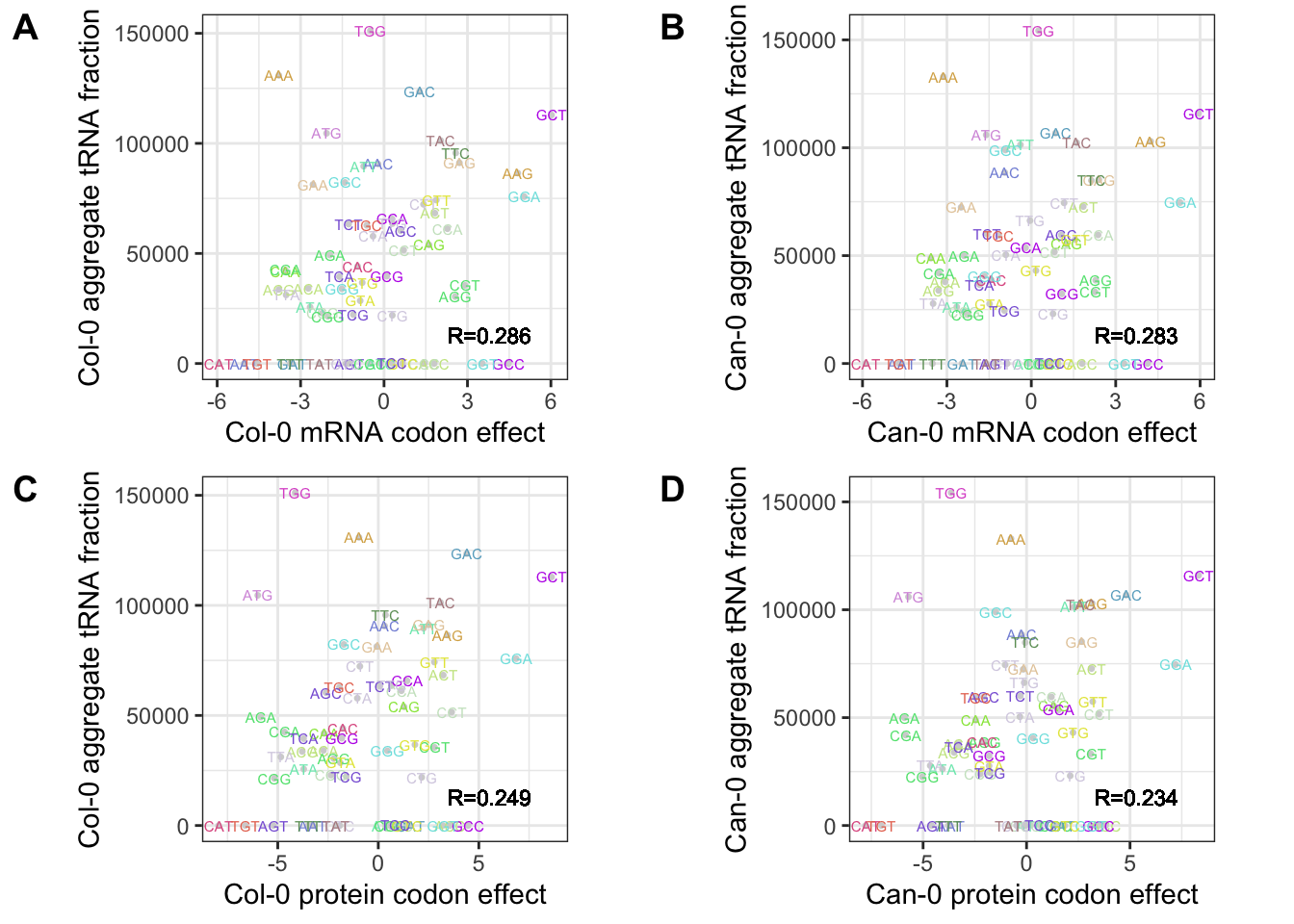
